# Contemporary and historical selection in Tasmanian devils (*Sarcophilus harrisii*) support novel, polygenic response to transmissible cancer

**DOI:** 10.1101/2020.08.07.241885

**Authors:** Amanda R. Stahlke, Brendan Epstein, Soraia Barbosa, Mark J. Margres, Austin Patton, Sarah A. Hendricks, Anne Veillet, Alexandra K Fraik, Barbara Schönfeld, Hamish I. McCallum, Rodrigo Hamede, Menna E. Jones, Andrew Storfer, Paul A. Hohenlohe

## Abstract

Tasmanian devils (*Sarcophilus harrisii*) are evolving in response to a unique transmissible cancer, devil facial tumour disease (DFTD), first described in 1996. Persistence of wild populations and the recent emergence of a second independently evolved transmissible cancer suggest that transmissible cancers may be a recurrent feature in devils. Here we compared signatures of selection across temporal scales to determine whether genes or gene pathways under contemporary selection (6-8 generations) have also been subject to historical selection (65-85 million years), and test for recurrent selection in devils. First, we used a targeted sequencing approach, RAD-capture, to identify genomic regions subject to rapid evolution in approximately 2,500 devils in six populations as DFTD spread across the species range. We documented genome-wide contemporary evolution, including 186 candidate genes related to cell cycling and immune response. Then we used a molecular evolution approach to identify historical positive selection in devils compared to other marsupials and found evidence of selection in 1,773 genes. However, we found limited overlap across time scales, with historical selection detected in only 16 contemporary candidate genes, and no overlap in enriched functional gene sets. Our results are consistent with a novel, multi-locus evolutionary response of devils to DFTD. Our results can inform management actions to conserve adaptive potential of devils by identifying high priority targets for genetic monitoring and maintenance of functional diversity in managed populations.

## Introduction

Species are subject to selection by pathogens throughout their evolutionary history, shaping lineage diversification and leading to complex cellular and molecular defensive mechanisms (1). Still, emerging infectious diseases (EIDs) can cause mass mortality and, given sufficient reproduction and genetic variation, initiate rapid adaptive evolution in a naïve host population (2). Although the prevalence and severity of EIDs in wildlife populations is now well-recognized (3–6), we are just beginning to understand the evolutionary impacts of disease in wildlife. We have a relatively short recorded history of infectious disease in wildlife, and therefore a limited ability to predict outcomes or intervene when warranted (7, 8).

High-throughput DNA-sequencing techniques and high-quality annotated reference genomes have revolutionized our ability to monitor and identify mechanisms of evolutionary responses to pathogens (8–10). Inter-specific comparisons of non-synonymous and synonymous variation (dN/dS) within protein-coding regions have long been used to identify positive selection at immune-related loci (11–13). At the population level, rapid evolution in response to disease can be detected by tracking changes in allele frequency before, during, and after the outbreak of disease (14, 15). Intra-specific comparisons across populations can reveal to what extent the evolutionary response to disease is constrained by limited genetic mechanisms or variation for adaptation (16). Reduced representation techniques such as restriction-site associated DNA-sequencing (RADseq) (17) have made the acquisition of genome-wide, time-series genetic data more accessible in non-model systems (18). By integrating these resources and tests of selection at differing temporal scales, we can assess whether species that show rapid evolution in response to contemporary pathogens also show evidence of historical selection to similar pathogens.

A striking example of an EID acting as an extreme selective force in wildlife is devil facial tumour disease (DFTD), a transmissible cancer first described in 1996 afflicting wild Tasmanian devils (*Sarcophilus harrisii*) (19). Tasmanian devils are the largest extant carnivorous marsupial, with contemporary wild populations restricted to the Australian island of Tasmania. As a transmissible cancer, DFTD tumour cells are transmitted between hosts, behaving as a pathogen (20). Transmission typically occurs as devils bite each other during the mating season after devils have reached sexual maturity (21, 22). With few notable exceptions documenting regression (23), DFTD tumors escape recognition, become malignant, and can kill their hosts within six months (24). Starting from a single Schwann cell origin (25), DFTD has now swept across nearly the entire species range (Figure 1A). Devil populations have declined species-wide ~80% (26) with local declines in excess of 90% (27). Nonetheless, population genomic studies have shown that devils are rapidly evolving in response to DFTD (2, 28, 29), and DFTD has been spontaneously cleared (i.e., regressed) in some individuals (23). Long-term field studies and simulation modelling have predicted that cyclical co-existence or DFTD extirpation are more likely scenarios than devil extinction (30). This is particularly alarming because devils have notoriously low genome-wide diversity, attributed to climate- and anthropogenic-induced bottlenecks (31–33). Depleted genetic diversity at immune-related loci has likely further contributed to DFTD vulnerability (34).

**Figure 1.**
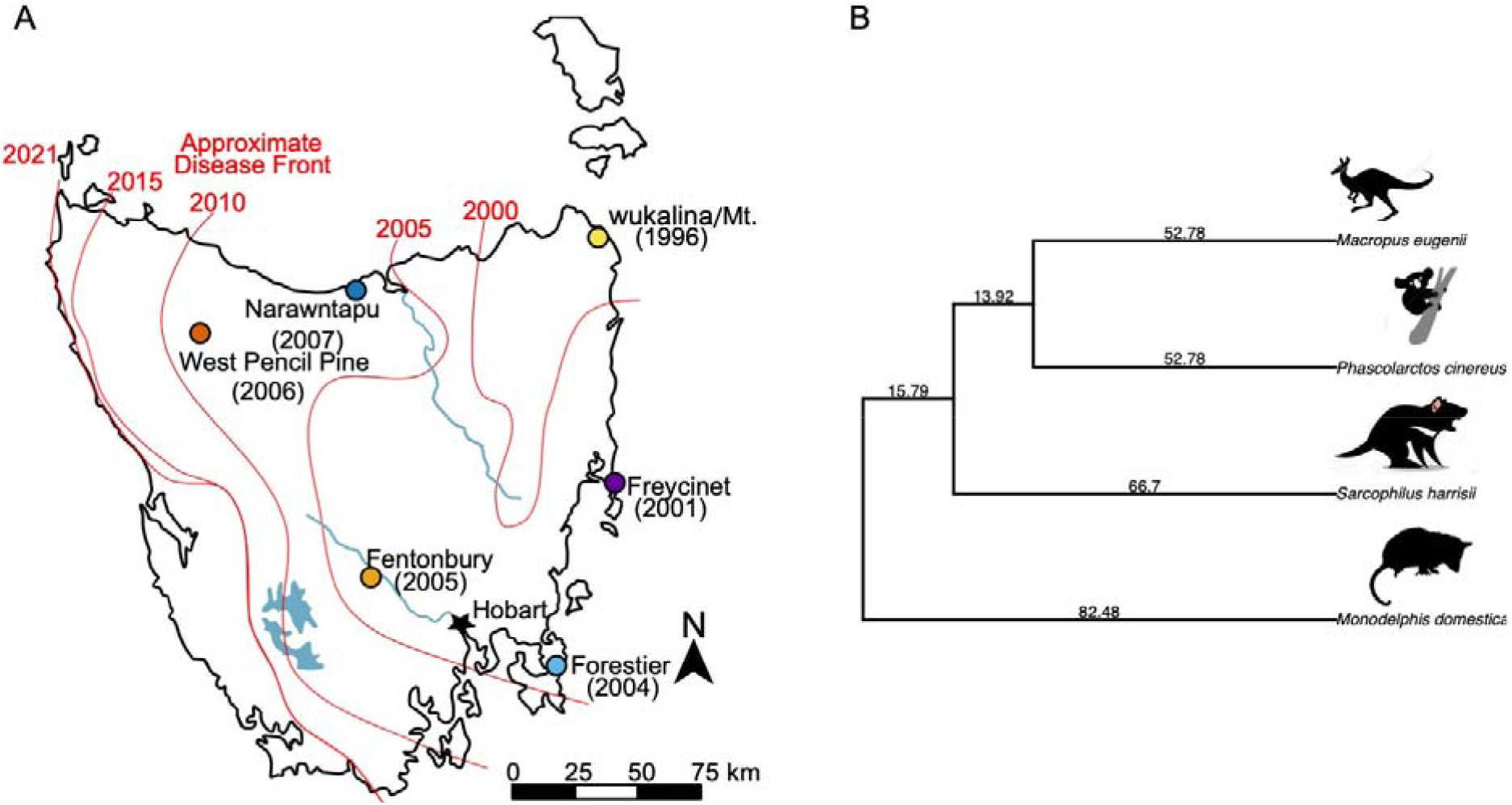
A) Map of the six contemporary sampling locations relative to disease prevalance over time (red lines) with the year of first detection labled at each site. B) Reduced, unrooted time-calibrated phylogeny (59) of marsupials used to estimate genome-wide historical selection on the devil lineage with estimated divergence times (Ma) indicted along edges. Devil cartoon by David Hamilton. Wallaby, koala, and opossum digital images retrieved from http://www.shutterstock.com/amplicon. From top to bottom: The tammar wallaby (*Notamacropus eugenii*), koala (*Phascolarctos cinereus*), Tasmanian devil (*Sarcophilus harrisii*), and South American grey-tailed opossum (*Monodelphis domestica*).

Despite transmissible cancers being exceedingly rare across animals, a second independent transmissible cancer in devils, DFT2, was described in 2014 (35, 36). Comparative and functional analyses of DFTD and DFT2 showed similar drivers of cancerous mutations and tissue type of origin (37). Low genetic diversity, chromosomal fragility (38), a reportedly high incidence of non-transmissible neoplasms (39), and injury-prone biting behaviour (40) may contribute to a predisposition to transmissible cancers in devils (41). These findings suggest that transmissible cancers may be a recurring selective force in the Tasmanian devil lineage. If so, this leads to the hypothesis that the genes and genetic pathways associated with the ongoing evolutionary response to DFTD may have experienced recurrent historical selection in the devil lineage from previous transmissible cancers.

Because of the threat of DFTD and DFT2 to devil populations, there are ongoing conservation efforts, including the establishment of a captive devil insurance meta-population. The insurance population is managed to maintain genome-wide genetic diversity and serve as a source for re-introductions in an effort to increase genetic diversity and size of wild populations (42). To inform conservation efforts, it is important to understand what types of genetic variation in natural populations may allow for evolutionary rescue from disease and maintain adaptive potential for future threats (43). Given evidence for rapid evolution in response to DFTD, monitoring of genetic variation at candidate adaptive loci could help evaluate adaptive potential of wild populations (43, 44). In heavily managed (e.g. captive) populations, loci associated with an adaptive response to disease could be included in genotyping panels for maintaining genetic diversity (45).

Here we identify targets of selection and signatures of adaptation at both contemporary (6-8 generations) and historical (65-85 million years) scales in Tasmanian devils. First we test for evidence of contemporary genomic response to selection by genotyping thousands of individuals sampled at several time points across six populations, using RAD-capture (46) to target nearly 16,000 loci (47). Next we identify signatures of historical selection in the devil lineage by comparing across marsupial species with annotated genomic sequence data. Then, we test for evidence of recurrent selection by examining shared contemporary and historical signatures of selection, in terms of either specific loci, genes or functional genetic pathways.

If transmissible cancer is a novel selective force acting on Tasmanian devils, we expect that genes under contemporary selection by DFTD will be different from those with a signature of historical positive selection. Alternatively, if transmissible cancer is a recurrent selective force in the devil lineage that targets the same set of genes repeatedly, we may expect a conserved response among populations and an overrepresentation of the same genes or pathways under both contemporary and historical selection. However, if there are multiple genetic pathways that could be involved in a response to recurrent transmissible cancers, we may expect a polygenic response across contemporary populations and little overlap between contemporary and historical timescales. These alternatives can inform conservation efforts to manage genetic diversity for resilience in natural devil populations, and any genes or functional pathways that show both contemporary and historical selection may be relevant to cancer resistance more broadly.

## Materials and Methods

### Contemporary Selection

We used the RAD-capture method (combining RADseq and sequence capture) (46) to conduct targeted genotyping of single-nucleotide polymorphisms (SNPs) across 2,562 unique individuals from multiple Tasmanian devil populations, sampled both before and after DFTD appeared in each population (Figure 1A, Table 1, Supplemental Table S1) (29, 48). We constructed RAD-capture libraries following Ali et al (2016), using the restriction enzyme PstI and a capture array targeting 15,898 RAD loci selected for membership in one of three functional categories: 1) those showing signatures of DFTD-related selection from previous work (2), 2) loci close to genes with known cancer or immune function, and 3) loci widely distributed across the genome (See 29, 47 for more details on the devopment of this array.). See Supplemental Materials S1 for multiplexing, read processing, and SNP genotyping details.

**Table 1.** Number of adults sampled before and after the year of first detection of DFTD at each site. See Supplemental Table S1 for sample size for each year at each locality.

| **Location** | **Year of First**<br>**Detection** | **Samples**<br>**Before** | **Samples**<br>**After** | **Total** |
| --- | --- | --- | --- | --- |
| wukalina/Mt. William | 1996 | 0 | 155 | 155 |
| Freycinet | 2001 | 300 | 382 | 682 |
| Forestier | 2004 | 131 | 552 | 683 |
| Fentonbury | 2005 | 99 | 169 | 268 |
| West Pencil Pine | 2006 | 52 | 348 | 400 |
| Narawntapu | 2007 | 224 | 150 | 374 |
| **Total** |  | 806 | 1756 | 2562 |

To account for the expected high rates of genetic drift within populations, we used a composite statistic to compare signatures of selection across populations. We identified candidate SNPs as the top 1% of a de-correlated composite of multiple signals score (DCMS) (49), which combined the results of three analyses: change in allele frequency in each population after DFTD (*Δaf*), and two methods that estimate strength of selection from allele frequencies at multiple time points in multiple populations, the method of Mathieson & McVean (14) (*mm*), which allows the estimated selection coefficient to vary over space; and spatpg (15), which allows the selection coefficient to vary over time and space. Individuals were assigned to generational-cohorts based on their estimated years of birth (Supplemental Table S1). We estimated *Δaf* for five locations at which we had sampling both before and after DFTD was prevalent, according to DNA collection date and estimated date of birth, combining multiple cohorts when applicable (Table 1; Supplemental Table S1). Both time-series methods (mm and spatpg) incorporate estimates of effective population size, which ranged from 26-37 according to a two-sample temporal method (50, 51) (Supplemental Table S3). DCMS reduces the signal-to-noise ratio by combining p-values from different tests at each SNP while accounting for genome-wide correlation among statistics. We included SNPs with results from at least eleven of the twelve individual tests (*Δaf* for five populations, *mm* for all six populations, and *spatpg*) and weighted based on the statistics with results at that SNP. To characterize the role of standing genetic variation in rapid evolution (52), we visualized the initial allele frequencies of each population for each analysis of contemporary evolution (Supplemental Figures S2, S9). We evaluated repeatability among populations by comparing population-specific p-values of *Δaf* and *mm* with the R package dgconstraint for a similarity index called the C-score, where 0 indicates no similarity between populations (16). See Supplemental Materials S1 for details of each analysis.

### Historical Selection

We combined existing genomic resources for the South American grey-tailed opossum (*Monodelphis domestica*) (53) and tammar wallaby (*Notamacropus eugenii*) (54) from the Ensembl database (55) and the recently published transcriptome assembly of the koala (*Phascolarctos cinereus*) (56) to identify genome-wide signatures of positive selection in devils, relative to these other species using the branch-site test of PAML (Phylogenetic Analysis by Maximum Likelihood) (57, 58). We compiled alignments of orthologous genes and reduced the marsupial time-calibrated phylogeny of Mitchell et al. (59) to those species for which annotated full genomes are available (Figure 1B). The branch-site test compares likelihood scores for two models which estimate dN/dS among site classes of a multi-sequence alignment, allowing dN/dS to exceed 1 (positive selection) in a proportion of sites along a single branch in the alternative model. We reduced the potential for false positives by filtering any putative orthologs with extreme sequence divergence (S > 2), measured as the sum of synonymous mutations per gene (S), and ensuring alignments of nucleotides were longer than 100 bp (60, 61). We identified historical candidates with the likelihood-ratio test, comparing the likelihoods of the alternative and neutral models with one degree of freedom and an α = 0.05. Historical candidates were those with estimates of dN/dS > 1 along the devil branch and FDR > 0.05 after correcting for multiple testing (62). See Supplemental Materials S1 for details regarding orthology identification and PAML implementation.

### Recurrent Selection

We refer to genes under both contemporary and historical selection as candidates for recurrent selection. To test whether genes under contemporary selection differed from genes under historical selection, we first tested for significant overlap with Fisher’s one-tailed test. To test for differences in the strength of selection, we compared the distributions of dN/dS and the proportion of sites per gene found under positive selection among candidates for recurrent selection to all other historical candidates from the genome-wide background using nonparametric tests of equality, the Kolmogorov-Smirnov test (63), which is more sensitive to the centre of the distributions, and the Anderson-Darling test (64), which is more sensitive to extreme values of the distribution and often has more power. To identify and compare key mechanisms of adaptation among candidate genes from each set, we used gene ontology (GO) term enrichment analysis using the SNP2GO package (65), the PANTHER web-interface (66), and in gene sets of the molecular signatures database (MsigDB), using the subset of genes tested for each test as the respective background set (67). We capitalized on the wealth of ongoing research in devils and DFTD by comparing our contemporary and historical candidates to those previously identified using different datasets and analytical approaches (2, 28, 29, 47, 68). See Supplemental Materials S1 for details of these comparisons.

## Results

### Genomic Data

To test for contemporary selection, we sampled a total of 2,562 individuals across six localities of Tasmania before and after DFTD prevalence (Table 1; Supplemental Table S1; Figure 1A), with a RAD-capture array (47). After filtering, we mapped a total of 517.7 million reads against targeted loci. The mean final coverage of targeted loci was 14.8x, with 76.6% of all samples having coverage of at least 5x (Supplemental Figure S1). After filtering, we retained 14,585 – 22,878 SNPs for downstream analysis, depending on the sampled time point and population.

### Evidence for contemporary selection

Among each elementary test for selection signatures, 161 – 232 SNPs (depending on population) were in the top 1% of allele shifts following disease (Δ*af*), 209 – 217 were in the top 1% of *mm* scores, and 213 were in the top 1% of spatpg scores (Supplemental Table S4, Figures S7–8). Across populations and elementary tests for contemporary selection (Δ*af, mm, spatpg*), *p*-values were not correlated (Pearson’s r < 0.155 for all tests; Supplemental Figure S10). The computed repeatability indexes for population-specific responses Δ*af* and *mm* were C_Δ*af*_ = 4.86 (p 1e-04) and C_*mm*_= 3.72 (p = 1e-04), which implies a low, but significant level of repeatability (16). In the top 1% of DCMS scores (≥ 1.167), we identified 144 candidate SNPs for contemporary selection by DFTD; of these, 79 had annotated genes (186 total) within 100 kb (Figure 2; Supplemental Table S5). The initial frequencies for candidate SNPs were not skewed toward intermediate frequencies (Supplemental Figure S9).

**Figure 2.**
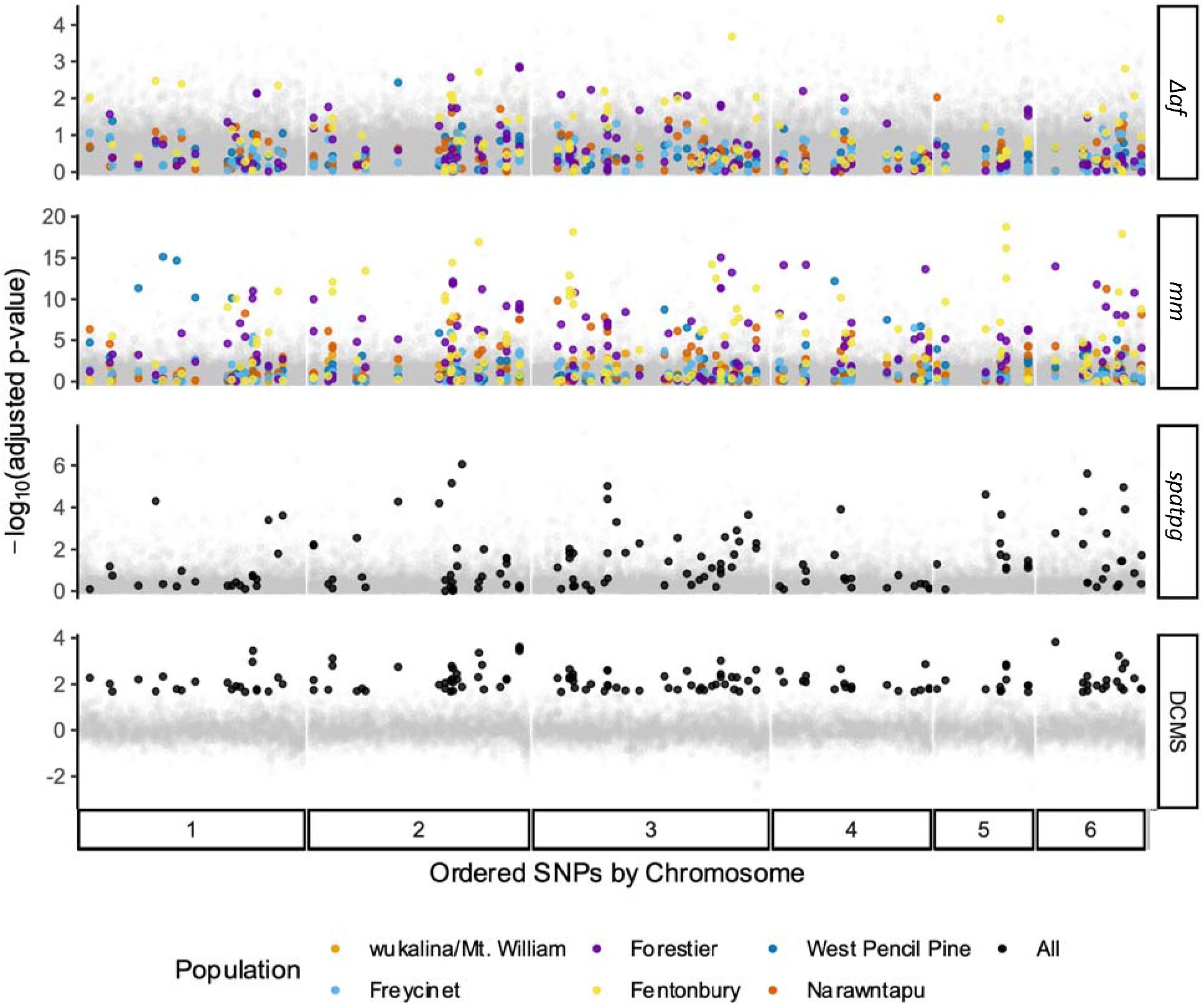
Results of each elementary test of contemporary selection across populations and the composite scores for final candidate (filled points) and noncandidate (opaque grey points) SNPs, ordered by chromosome and colored by population when applicable. From top to bottom: Change in allele frequency (Δ*af*), Mathieson and McVean (*mm*) (14), *spatpg* (15), de-correlated composite of multiple signals (*DCMS*) (49).

Comparing our contemporary candidates and those previously identified in devils with selection and genome-wide association analyses (28, 29, 47, 68), we found many overlapping genes (discussed below). Notably, we found significant enrichment of candidates previously associated with DFTD-related phenotypes in females (14 genes, *p* = 4.2e-08, Odds ratio=7.3) (47). Gene ontology enrichment analysis found middle ear morphogenesis (GO:0042474) significantly enriched among contemporary candidate SNPs (FDR < 0.05). Five candidate SNPs were within the 100 kb window of two genes associated with this term: EYA1 and PRKRA. Both EYA1 and PRKRA are involved in cell proliferation and migration and implicated in tumour suppression and angiogenesis (69–71).

### Evidence for historical selection

Of the 18,788 genes annotated in the devil reference genome, 6,193 had 1-to-1 orthologs in at least three of the four marsupial genomes and an appropriate sequence divergence (S<2). Using the branch-site test for positive selection in PAML, we found a total of 1,773 genes to be candidates for historical positive selection (Supplementary Table S6). Estimates of dN/dS spanned the full range of possible values, from 1.05 – 999 and proportion of sites with substitutions per gene ranged from 0.01 – 0.78 (Fig. 3). The majority of genes were classified as having a molecular function of binding (GO:0005488) or catalytic activity (GO:0003824); a plurality involved in cellular processes (GO:0009987) or biological regulation (GO:0065007); and a plurality as participating in the Wnt signalling pathway (P00057). None of these pathway classifications were significantly enriched.

**Figure 3.**
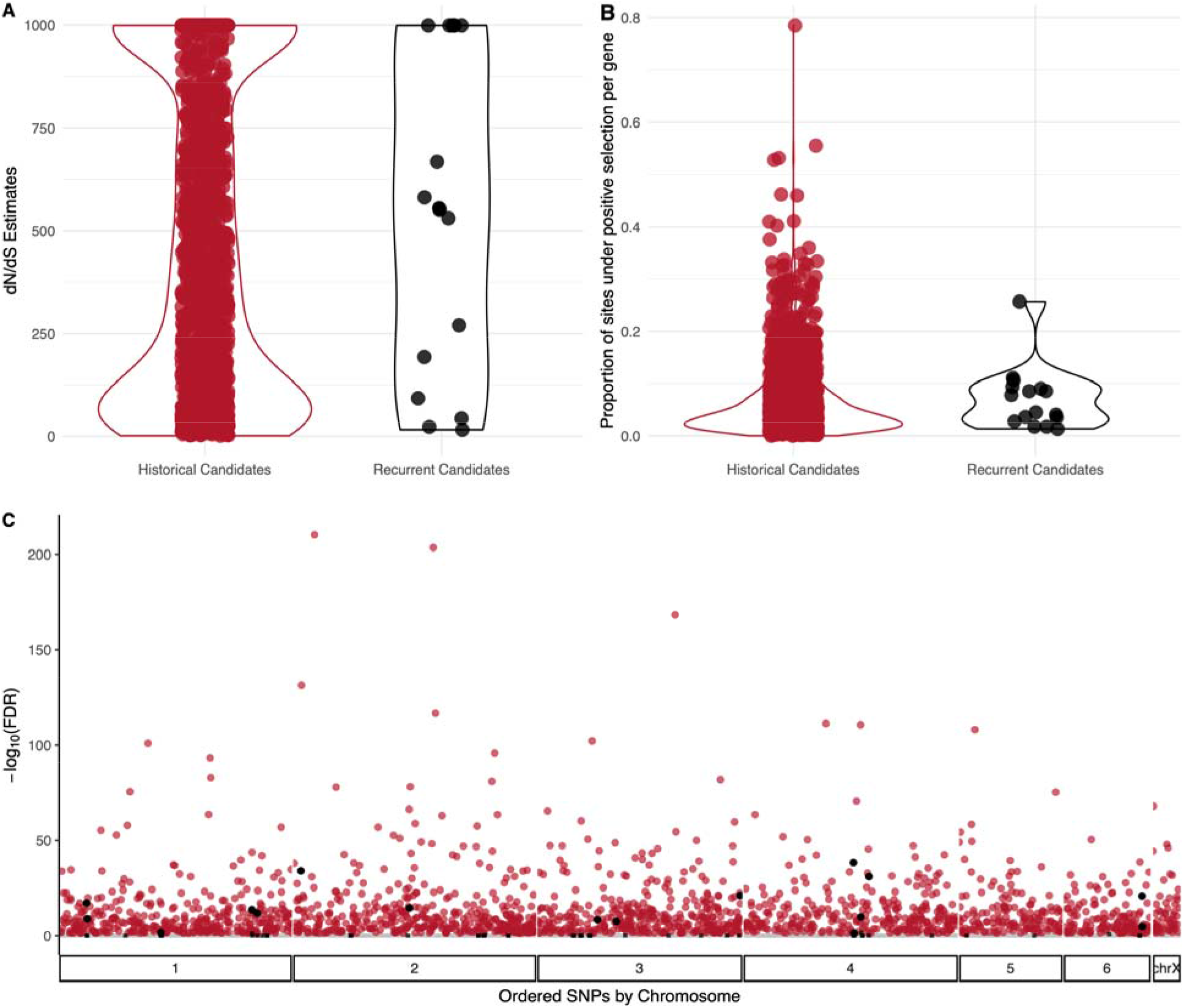
A) Estimates of dN/dS and B) the proportion of sites under positive selection (57) for historical candidates across the genome-wide background (red; N=1,982) and candidates for recurrent selection (genes with significant results for both historical and contemporary selection; black). Each point represents the respective result for a single gene. C) Distribution of -log(FDR) for historical selection across all 6,193 genes tested (gray squares, non-significant at both scales; black squares, significant contemporary and non-significant historical; red circles, non-significant contemporary and significant historical; black circles, significant at both scales).

### Recurrent selection

Of the 186 contemporary candidate genes, 68 had 1-to-1 orthologs among the four marsupials and were tested for historical selection. Sixteen genes showed evidence of historical selection and are thus candidates for recurrent selection (dN/dS > 1, FDR < 0.05; Supplemental Table S6). Contemporary candidates were not enriched for historical selection according to Fisher’s test (Odds ratio = 0.0, *p* = 1). Among the 16 recurrent candidates, dN/dS estimates spanned 15.7 – 999 and proportion of sites per gene 0.01 – 0.25 (Fig. 3). According to the Anderson-Darling and Kolmogorov-Smirnov tests of equality, neither distributions of dN/dS estimates (Fig. 3a; A.D. *p* = 0.86; K.S. *p* = 0.58), nor proportion of sites (Fig. 3b; A.D. *p* = 0.49; K.S. *p* = 0.48) differed between candidates for recurrent selection (in black) and historical candidates (in red).

After correcting gene set enrichment for multiple testing (FDR < 0.05) and requiring at least 10 genes in the background set, we did not find functional enrichment of any MSigDB gene sets among recurrent candidates or shared between both contemporary and historical sets. Importantly, the permutation test of shared gene sets found *fewer* shared between historical and contemporary selection than expected by chance (*p* < 0.001, Supplemental Figure S11).

## Discussion

### Contemporary Responses to DFTD

Using a targeted set of nearly 16,000 loci, we detected widespread evidence of a response to selection by DFTD across the Tasmanian devil genome. Our results extend previous work that has shown genomic evidence of a response to DFTD in wild populations (2, 28, 29, 72). Here we greatly increased the sample size of individuals and genetically independent populations for greater power, resulting in strong evidence of a response to selection widely distributed across the genome. We found greater similarity across populations within analytical approaches than among methods within populations and relatively low, but significant repeatability across populations. This result is consistent with rapid, polygenic evolution facilitated by selection for standing variation within populations that was present prior to disease arrival. This timescale (3 - 8 generations) would likely be too short for new mutations or migration to play a substantial role in DFTD response, and genetic variation is shared across the species range, despite geographic population structure (29, 73).

In line with previous population genomic studies (2, 28, 29, 47, 68), our analysis of contemporary evolution detected a putatively adaptive response related to the immune system, cell adhesion, and cell-cycle regulation (Supplemental Table S5). Our GO enrichment result for middle ear development (GO:0042474) among contemporary candidates may highlight selection for interactions with the peripheral nervous system and cell proliferation. Genes annotated with nervous system associations may indicate selection for behavioural changes (28), or highlight importance and vulnerability of peripheral nerve repair by Schwann cells in devils, given the prevalence of biting and Schwann cell origin of DFT (41). Significant overlap for genes associated with devil infection status (case-control), age, and survival (47) among our contemporary candidates is a strong indicator that these contemporary candidates likely confer relevant phenotypic change. We also confirmed five (CRBN, ENSSHAG00000007088, THY1, USP2, C1QTNF5) of seven candidates identified previously (2) in a genome scan for loci under selection from DFTD in three of the same populations (Freycinet, Narawntapu, and West Pencil Pine). In contrast, we identified those five and only two more (TRNT1 and FSHB) of 148 candidates from a re-analysis of that same dataset which studied population-specific responses (28).

Among genes that have been associated with host variation responsible for tumour regression on devils (68, 74), we found only JAKMIP3, a Janus kinase and microtubule binding protein (74), in our list of contemporary candidates. However, we found devil regression candidates TL11, NGFR, and PAX3, which encodes a transcription factor associated with angiogenesis (75); as well as GAD2, MYO3A, and unannotated ENSSHAG00000009195 (74), among population-specific candidates for allele frequency change (Δ*af*), possibly reflecting differences in test sensitivities. Overall, the paucity of candidates shared between our contemporary analysis and regression studies suggests that regression may not be the dominant form of phenotypic response to DFTD; to date tumour regression has only been detected in a few populations (74) not represented in our study.

### Historical selection in the devil lineage

With our genome-wide molecular evolution approach (57), we found widespread historical positive selection across the devil genome in about 28% of all 6,249 orthologs tested (Supplemental Table S6). The branch-site test is known to be less conservative than related models, particularly when divergence among species is large (76), but the rates of historical selection we found in devils are similar to those described in other taxa; e.g. 23% of genome-wide orthologs among 39 avian species using a similar approach (13).

We did not find preferential positive selection for immunity-related genes, as has been shown in primates (1), eutherian mammals more generally (77), and birds (13). Instead, we found the highest proportion of pathways under historical selection to be functionally classified within the Wnt pathway, a signalling cascade regulating cell adhesion and implicated in carcinogenesis (78). As genomic resources grow and improve in marsupials (10), interspecific analyses for positive selection at finer scales may reveal more recent and specific selection targets in Tasmanian devils. Our ability to detect historical selection due to transmissible cancer in devils could be improved by genome assembly efforts among more closely-related Dasyuridae, as well as complementary annotation.

### Comparing Contemporary and Historical Timescales

Remarkably few transmissible cancers have been discovered in nature (79, 80), and yet two of those independent clonally-transmitted cancers have been discovered in Tasmanian devils in less than 20 years. This and the observed rapid evolutionary response to disease suggest that transmissible cancers may be a recurrent event in devils. We found no significant overlap of historical and contemporary selection at either individual genes or functional gene sets. This does not rule out the possibility of prior transmissible cancers in devils; but it suggests that if transmissible cancers have been a recurrent feature of devil evolution prior to DFTD, they did not generally impose selection on the same set of genes or genetic pathways that show a contemporary response to DFTD. Nonetheless, the 16 candidate genes showing both historical and contemporary evidence for selection (Supplemental Table S7) raise interesting targets for understanding adaptively important variation in devils.

The 16 candidate genes for recurrent selection (Supplemental Table S7) are generally related to three main themes: transcription regulation, the nervous system, and the centrosome. Four of these candidates for recurrent selection were previously associated with disease-related phenotypes (47). We additionally found 82 historical candidates previously identified in the top 1% of SNPs associated with disease-related phenotypes with three represented in the top 0.1% associated with large-effect sizes for female case-control and survival (47). This overlap lends support to the hypothesis of recurrent selection by transmissible cancers, but was not significant (*p* = 1, odds ratio= 0). Both our contemporary selection analysis and the genome-wide association study (GWAS) approach used by Margres and colleagues (47) are statistically limited by small populations, sample size, and the time scale over which DFTD-related selection has occurred. By considering the complement of these results together, the overlapping historical, GWAS, and contemporary candidates may still be targets of recurrent selection along similar functional axes, potentially including transmissible cancer.

The low prevalence of candidates for recurrent selection and lack of shared functional gene set enrichment between both contemporary and historical signatures of selection suggest a novel response to DFTD compared to historic selection in the devil lineage. However, there are alternative hypotheses. For example, there could be redundancy in genetic mechanisms underlying resistance to transmissible cancers, potentially as a result of repeated selection for resistance, allowing selection to act across many loci (81). That is, the low genetic diversity observed in devils could be the result of widespread historical purifying selection resulting from transmissible cancers or other diseases (82), or historical bottlenecks due to climate change and habitat loss (31–33), that prevent a response to selection under DFTD at loci that are still associated with disease phenotypes.

The widespread contemporary evolution we found in devils reflects the recent prediction (83) that response to an emergent disease is most likely controlled by many genes conferring quantitative resistance (84), for example by reducing the within-host growth rate of tumors. DFTD is predicted to become less virulent in the short-term (30, 85). If DFTD persists long-term in the devil population with ongoing coevolution, it may lead to diversifying selection for specific, qualitative host resistance mechanisms (83). Indeed, phylodynamic analysis of DFTD as it spread across Tasmania supports the hypothesis that devils may be mounting a response; transmission rates have decayed such that DFTD appears to be shifting from emergence to endemism (85). Although host-genomic variation was not jointly considered in that study, the combined evidence of multiple studies demonstrating rapid evolutionary response of devils to DFTD, including this one, support these interpretations.

### Conservation Implications

Calls have been made to consider the historical context of adaptation when proposing conservation management solutions based on genomic results (86). Our analysis of historical selection largely supports the hypothesis that DFTD is a newly emerging and novel selective force, distinctly shaping today’s remaining wild devils. The targets of novel selection that we identified (Fig. 2, Supp Table S4) and their functional roles should be considered for prioritization of monitoring and conservation in light of DFTD. At the same time, the wide distribution of contemporary candidates across the genome also highlights the importance of standing genetic variation to continue to respond to unique selective forces, including local environmental factors (29). Genomic monitoring could be useful for maintaining both functional diversity at candidate loci and genome-wide variation in captive populations (45, 87, 88) and in the wild. Multiple genomic tools are available for targeted monitoring of large sets of loci (e.g. 89, 90) and could be used to track adaptive evolution and potential in the form of genetic diversity (43). However, before management decisions are made for specific genes, further work would need to identify favoured alleles and fitness effects for the genes we identified (Supplemental Table S5).

DFTD has yet to reach devils in the far west (Fig. 1a) and continues to circulate throughout the island. To maintain long-term adaptive capacity in the face of similar recurrent selective forces including DFT2 and potential future transmissible cancers, our results warrant (1) the monitoring of genetic variation in broad functional groups and (2) management strategies to maintain genetic diversity across those broad groups. Although these populations were not subject to DFT2 at the time of writing, an important and interesting future direction should examine the evolutionary response to DFT2 and could compare loci under selection by the two independent transmissible cancers. This study could provide a list of candidate loci for development of a genotyping panel for either purpose, with flexibility to target many or fewer loci. At the same time, given urgent and unpredictable present-day threats including not just emerging diseases but environmental change and population fragmentation, it is important that monitoring and population management also focus on maintaining genetic variation across the genome.

## Conclusion

Our results suggest that the contemporary evolutionary response to DFTD is mostly novel compared to the genome-wide signature of historical selection. Comparing the degree of overlap and distributions among contemporary and historical candidates did not support recurrent selection on a common set of genes in response to transmissible cancer. Our work contributes to mounting evidence of possible mechanisms by which devil populations are persisting and rapidly evolving in the face of DFTD despite overall low genetic diversity and population bottlenecks (2, 23, 47, 72, 91). Broadly, this type of approach can be applied to analyses of novel threats in wildlife populations in the current era of anthropogenic global change to guide monitoring and management actions focused on genetic adaptive potential.

## Data and Script Accessibility

Demultiplexed sequence data has been deposited at NCBI under Bio-Project PRJNA306495 (http://www.ncbi.nlm.nih.gov/bioproject/?term=PRJNA306495) and BioProject PRJNA634071 (http://www.ncbi.nlm.nih.gov/bioproject/?term=PRJNA634071). Code and tabular results are available at https://github.com/Astahlke/contemporary_historical_sel_devils and on Dryad.

## Acknowledgements

The authors would like to acknowledge Michael R. Miller, Sean M. O’Rourke, Cody G. Wiench, for their contributions to the RAD capture data; and Rebecca Johnson and Denis O’Meally for providing the koala transcriptome data. This work was funded by NSF grant DEB-1316549 and NIH grant R01-GM126563 to A.S., P.A.H., M.E.J., and H.I.M. as part of the joint NSF-NIH-USDA Ecology and Evolution of Infectious Diseases program. Genomics and bioinformatics were supported by an Institutional Development Award (IDeA) from the National Institute of General Medical Sciences of the NIH under grant number P30 GM103324. This work was supported by the Bioinformatics and Computational Biology Program at the University of Idaho in partnership with IBEST (the Institute for Bioinformatics and Evolutionary Studies). Sample collection was funded under Australian Research Council grants DE170101116, DP110102656, LP0989613 and LP0561120 to R.H., M.E.J., H.I.M., and A.S., an ARC Future Fellowship FT100100031 to M.E.J., and multiple awards from the University of Tasmania Foundation Eric Guiler Research Grant.

## Authors’ contributions

ARS, HIM, MEJ, AS, and PAH conceived and designed the study. RH and MEJ conducted fieldwork and sampling. ARS, BE, SB, SAH, AV, BS conducted genomics labwork. ARS, BE, SB, AP, SAH, AKF, and PAH conducted bioinformatic analyses. ARS wrote the manuscript with contributions from all authors.

## Supplementary materials S1

### Methods

#### Rapture Sequencing for Contemporary Selection

We multiplexed 672-868 individuals per lane of Illumina NextSeq, obtaining 2.4 billion 150 base pair (bp) paired-end reads. We later re-sequenced 2,379 individuals from those Rapture libraries on an Illumina HiSeq 4000 to increase coverage and confidence in genotype inference. Finally, we also incorporated data from standard RAD sequencing libraries on four additional individuals from two of the six populations, West Pencil Pine and Fentonbury (see 1, 2 for details.).

For each individual, we merged all available reads from across sequencing efforts. Then reads were de-multiplexed and low-quality reads were removed using process_radtags in Stacks using the ‘--bestrad’ option which checks for the single restriction enzyme cut site on either read; this step also removed any reads without recognizable barcodes or cut sites (3). The Stacks clonefilter program was used to remove potential PCR duplicates (4). Using bowtie2 (5), reads were aligned to the *S. harrisii* reference genome Devil_ref v7.0 (6), downloaded from Ensembl in June 2014. Population allele frequencies can often be estimated with greater accuracy and reduced bias than individual genotypes (7). To do this, we used ANGSD v0.910 (8). For each set of individuals, we calculated genotype likelihoods and estimated allele frequencies within regions using the settings in Supplemental Table S2. Regions on the X chromosome were excluded.

#### Spatial and temporal analysis of contemporary selection

First, we calculated allele frequency change after DFTD infection. Within the five locations for which we had sampling both before and after DFTD appearance, we estimated the magnitude and direction of allele frequency change at SNPs with data from at least 10 individuals at both time points and a minor allele frequency (MAF) ≥ 0.05 and a likelihood-ratio test p-value (for presence of a SNP from ANGSD) ≤ 10^−6^ in at least one of the time points. For this analysis, individuals were assigned to “before” or “after” time points based on the date of DNA collection. Table 1 presents the first year of DFTD detection in each population and samples collected from individuals after those years were considered “after.” DFTD could have been present at very low frequency prior to detection in some of these populations, but we believe using these dates of detection still provides a good estimate of pre-DFTD allele frequencies. We performed an arcsine (Fisher’s angular) transformation on the estimated allele frequencies to reduce bias induced by the allele frequency spectrum (9). The SNPs were ranked by the magnitude of change, and the fractional rank was used as a pseudo- p-value for the composite statistic (described in the Main Text).

Then, we identified SNPs with estimates of strong selection using two time-series approaches which account for a population structure: the method of Mathieson & McVean (10) (hereafter *mm*), which allows the estimated selection coefficient to vary over space; and *spatpg*, which allows the selection coefficient to vary over time as well as space (11). Individuals were divided into two-year cohorts based on their year of birth, starting with 1997 and ending with 2012 (Supplemental Table S1). Only SNPs with MAF ≥ 0.05, minor allele count ≥ 3, and p-value from ANGSD ≤ 10^−6^ in at least five population / cohort combinations were tested. For *mm*, we assumed the same effective population sizes as Epstein et al. (2016), and otherwise assumed similar effective population sizes where previous estimates did not exist (Supplemental Table S3). For *spatpg*, variance effective population sizes were estimated within the program with a bounded prior between 25 and 40. We created input allele count files for both *spatpg* and *mm* by multiplying allele frequencies estimated in ANGSD and the number of individuals and rounded to the nearest whole number. Due to computational limits, the dataset was randomly divided into 18 separate *spatpg* runs. Following *spatpg* recommendations in the manual (11), we calculated a support value for each SNP by taking the proportion of the posterior distribution (i.e. proportion of MCMC steps) for which the regression coefficient *β*, was non-zero (0 < *β* or 0 > *β*, whichever was smaller), where *β* describes the association between allele change and presence/absence of DFTD within a population. We multiplied these support values by two and treated them as pseudo- p-values when calculating a genomic inflation factor and the composite statistic.

Following the recommendations in Francois and colleagues (12), we adjusted the p-values of each test to reduce false positives. If the distribution of p-values was not uniform, we divided the Z-scores by the inflation factor. The inflation factor was calculated as the ratio between the median Z-scores and the expected median Z-scores for a *χ*^2^ distribution with one degree of freedom. The genomic inflation factor varied from 0.25 – 0.58 for *mm* analyses of each population. For *spatpg*, we found an inflation factor of 0.44.

Using the adjusted p-values and pseudo p-values from the individual analyses, we calculated the DCMS statistic (13). This statistic combines the p-values from different tests at each SNP while accounting for genome-wide correlation among statistics. For each SNP, we used a weight based on only the statistics with results at that SNP, and we only included SNPs with results from at least eleven of the twelve individual tests (Δ*af* for five populations, *mm* for all six populations, and *spatpg*).

We then divided DCMS by the number of defined tests to get a mean composite score and ranked SNPs by this mean score. Because DCMS is not defined when one of the p-values is one or zero, we replaced p-values of one with 0.99999, and values of zero (only occurred for spatpg) with 0.00005 before performing the calculation. In accordance with previous linkage disequilibrium estimates (2), we identified candidate genes within 100 kb of top SNPs, using bedtools (version 2.26.0) (14); and supplemented annotations of novel genes with the Ensembl Compara gene family pipeline (15).

#### Historical Selection

To test for historical selection we used the branch-site test of PAML (Phylogenetic Analysis by Maximum Likelihood; 16, 17), implemented in the Bio.Phylo toolkit (18) of BioPython (19). The branch-site test estimates the ratio of nonsynonymous-synonymous mutation rates (dN/dS) among aligned codons using a phylogenetic tree to allow for the appropriate evolutionary model to be employed. In the neutral model, all site classes and branches are constrained to dN/dS ≤ 1. In the alternative model, dN/dS of Site Class 2 is allowed to exceed 1 for only the foreground branch, while constraining the background branches to dN/dS ≤ 1.

The devil, wallaby, and opossum orthologous genes and respective sequences were mined from the Ensembl database with BioMart (20, 21). For the koala, we used the orthologs identified with blastx version 2.2.27+ (22) supplied by Johnson and colleagues (23). If splicing variants were available, only the first (most common) variant was retained for downstream data preparation. Only 1-to-1 orthologs were retained (i.e., paralogs were excluded). Orthologous gene tables were then reduced to genes with at least three of four possible sequences present and the respective species were pruned from the greater phylogeny. For the koala, we used an open reading frame finder, getorf from EMBOSS, (24) to generate amino acid sequences. The peptide sequence alignments were generated with MUSCLE version 3.8.31 (25, 26), then used to guide alignments of nucleotides with tranalign.

#### Functional Enrichment of Genes Under Selection

For contemporary candidate SNPs, we used the SNP2GO package (27) in the R environment (version 3.4.3). We filtered the most recent Gene Transfer File (Sarcophilus_harrisii.DEVIL7.0.100) and Ensembl gene ID GO term associations downloaded from Ensembl May 7, 2020 to include only genes which were within 100 kb of targeted loci and account for the biased subset of targeted genes. We limited enrichment analysis to GO terms with a minimum of one association in the reference set and allowed an extension window of 100 kb from a candidate SNP. For GO term enrichment analysis of historical selection, we used the PANTHER web-interface with HUGO gene names (28) and the reference set defined as all genes that were tested in PAML (29).

We only compared the distributions among genes for which dN/dS > 1 and were statistically significant according to the likelihood ratio test. To account for the bias induced by targeted sequencing among contemporary candidates, we defined the reference set as all genes that could have been detected in both tests for a given overlapping or enrichment analysis We capitalized on the wealth of ongoing research in devils and DFTD by comparing our contemporary and historical candidates to those previously identified using different datasets and analytical approaches (2, 30–33) using Fisher’s Exact Test implemented in the R package GeneOverlap (34). We then tested for overrepresentation of contemporary and historical candidates in gene sets of the molecular signatures database (MsigDB) (35). MSigDB contains several libraries of gene sets which allowed us to gain further insight to pathways that may be under selection in devils. We compared our contemporary and historical candidate gene lists to gene sets from the MsigDB Hallmark, Curated, Computational, Oncogenic Signatures, and Immunologic Signatures (36, 37). We built 2×2 contingency tables for each set of genes under positive selection and in each of the tested gene sets of MsigDB. Despite limitations, this overrepresentation method is straight-forward and flexible for non-model organisms and targeted sequencing. To identify intersecting gene sets, we first converted all Ensembl gene IDs of interest to HUGO annotations with Biomart. Second, we created appropriate background sets (of length N) by intersecting each MsigDB gene set with the respective list of all genes that were tested for selection. Lastly, from these contingency tables we computed overlaps, using the hypergeometric distribution (dhyper(c(0:x), m, n, x, log = FALSE), where c(0:x) is a vector of quantiles representing the number of genes both under selection and found in an MsigDB gene set; m is the number of genes in the candidate gene list, n is the number of genes in the candidate gene list but not in the MSigDB gene set, and x is the number of candidate genes in the MsigDB gene set list. After accounting for multiple testing with the Benjamini-Hochberg correction (38), we considered the overrepresentation result statistically significant with adjusted p-values < 0.05 and background gene sets greater than or equal to ten genes. Finally, we performed a permutation test to establish a null expectation for the rate of shared gene sets between contemporary and historical selection and compared the resultant empirical null to our observed proportion of shared gene sets. To do this, we randomly selected the same number of candidate genes with HUGO annotations (112) from the list of all HUGO annotated contemporary candidates (3,920) 1,000 times, with replacement, and performed gene set overlap analysis as above with only the known gene sets significantly overlapping with the historical candidates.

---

**Supplemental Figure S1.**
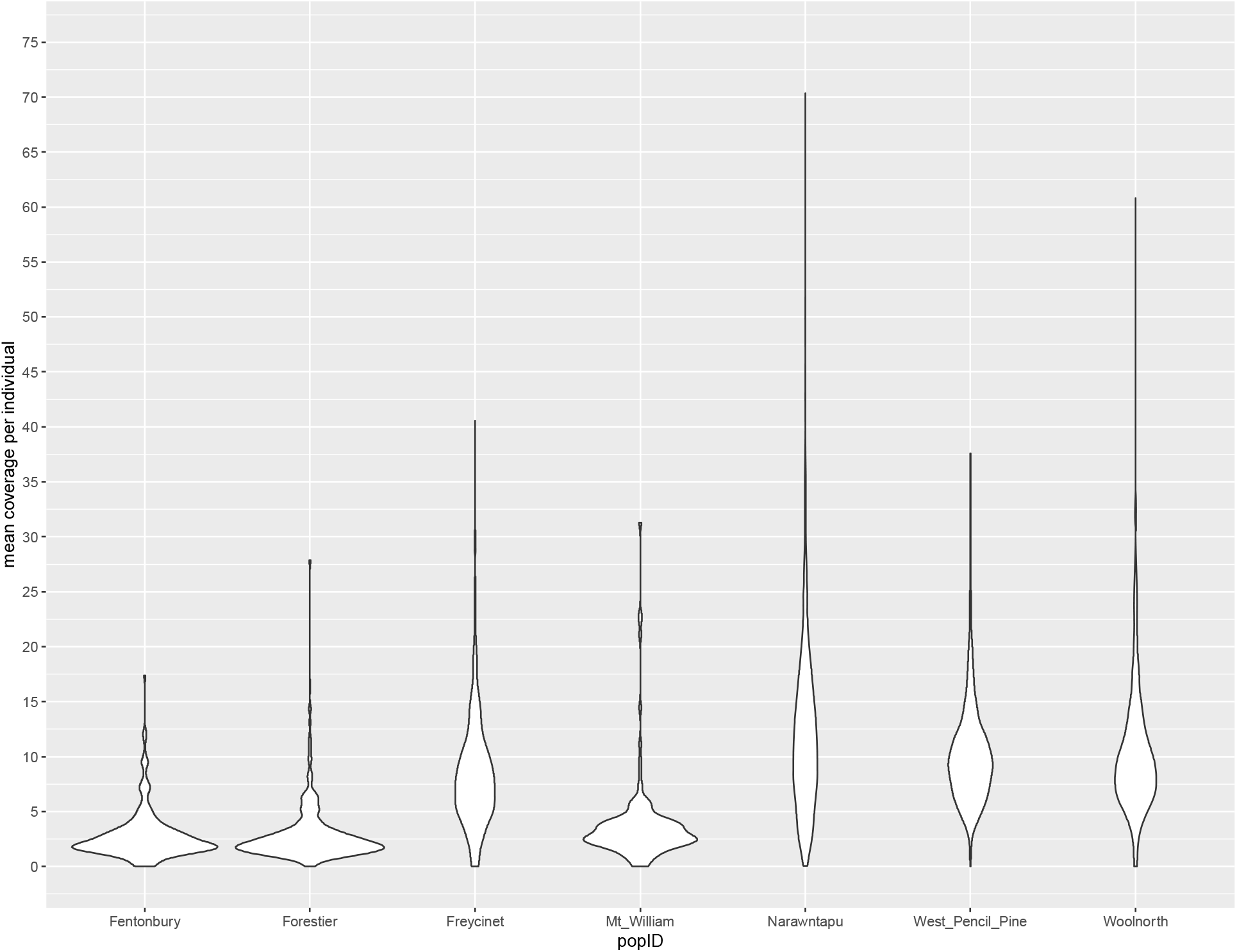
Mean coverage of individuals across populations at targeted loci

**Figure S2.**
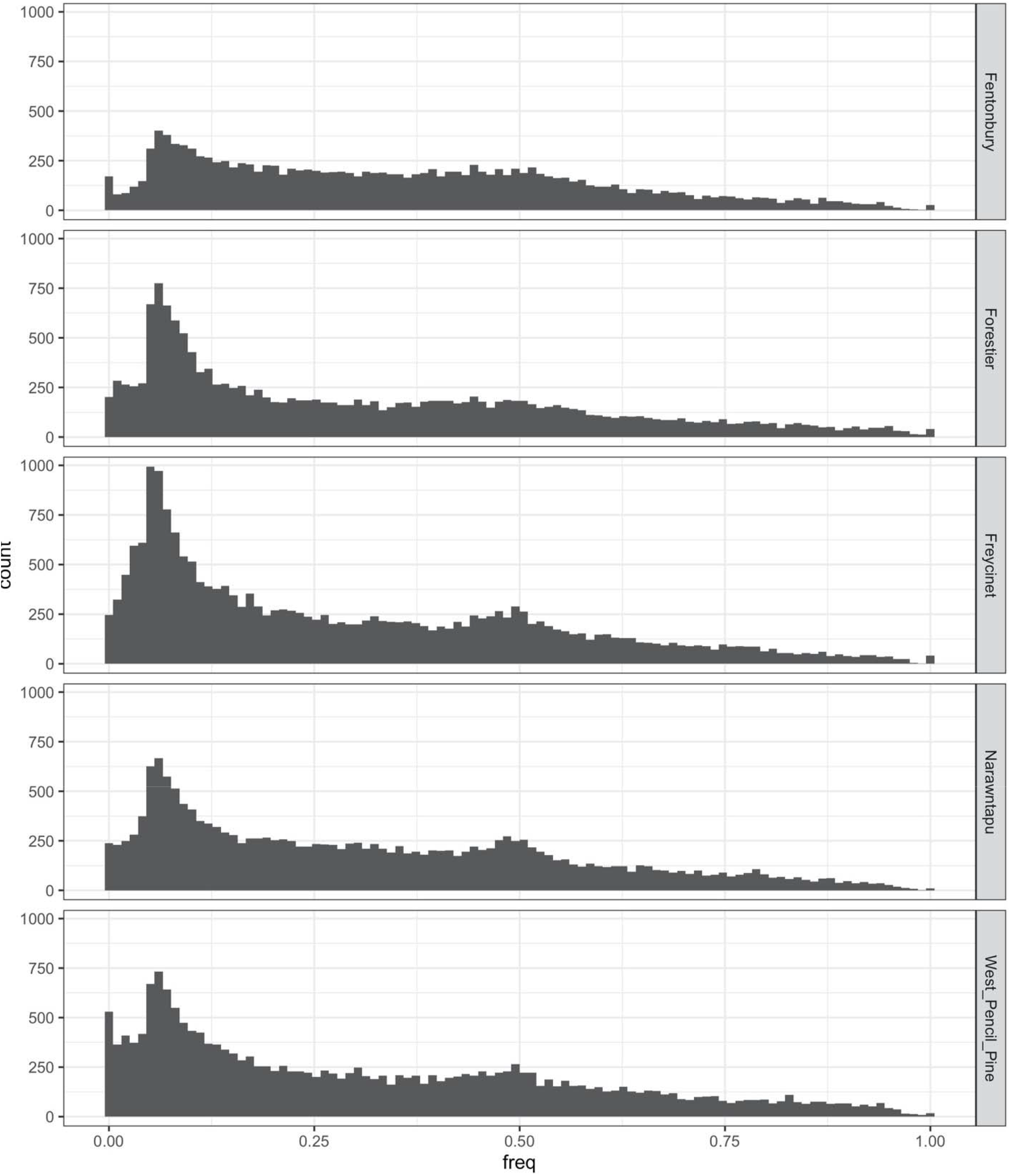
The folded allele frequency spectra for each population before DFTD became prevalent. wuaklina/Mt. William is not presented because it was first sampled in 2004, eight years after DFTD was first described at that locality.

**Figure S3.**
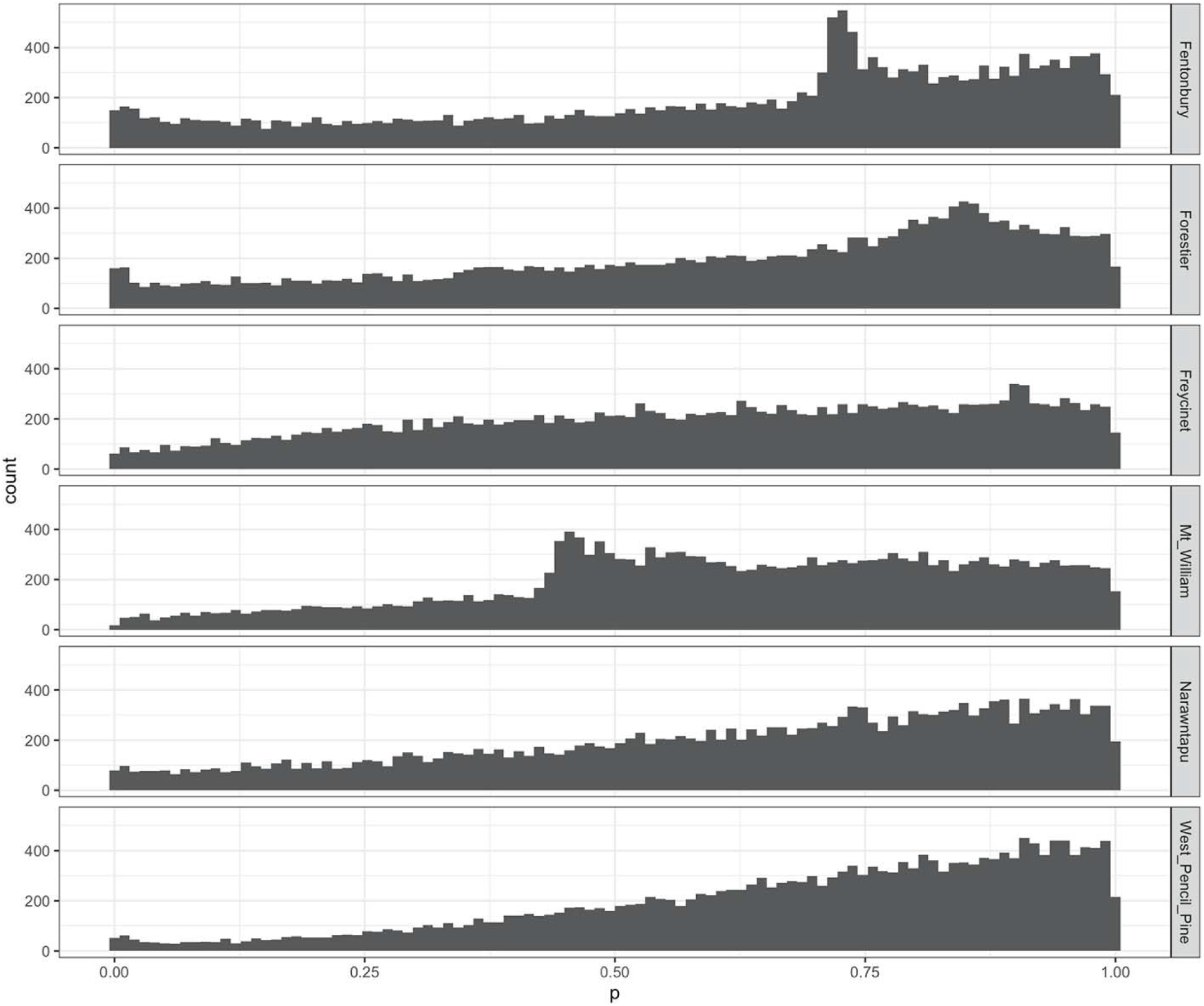
Un-adjusted p-values for all SNPs of each population analysed with *mm* (10).

**Figure S4.**
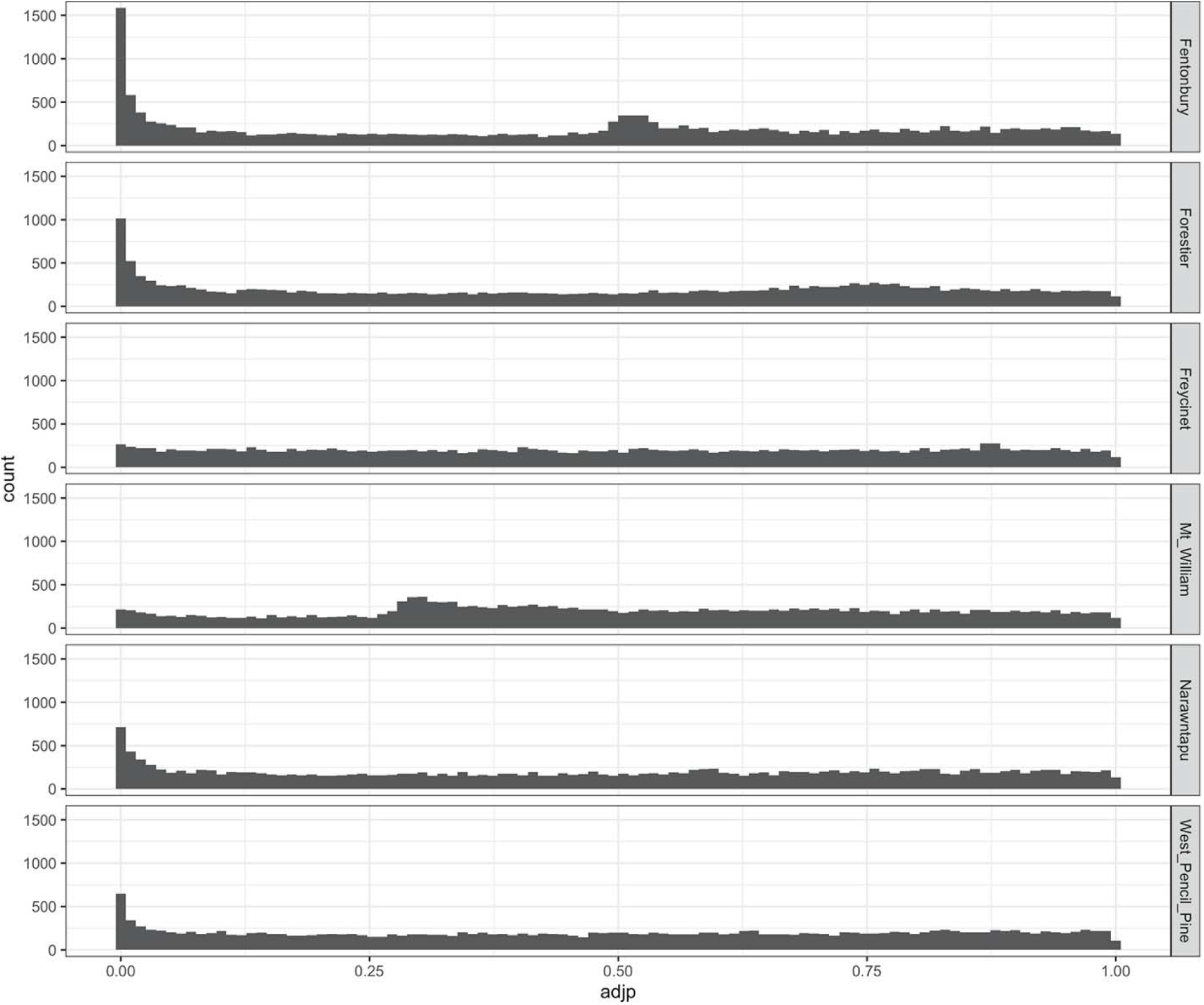
Adjusted p-values (12) for all SNPs of each population analysed with *mm* (10).

**Figure S5.**
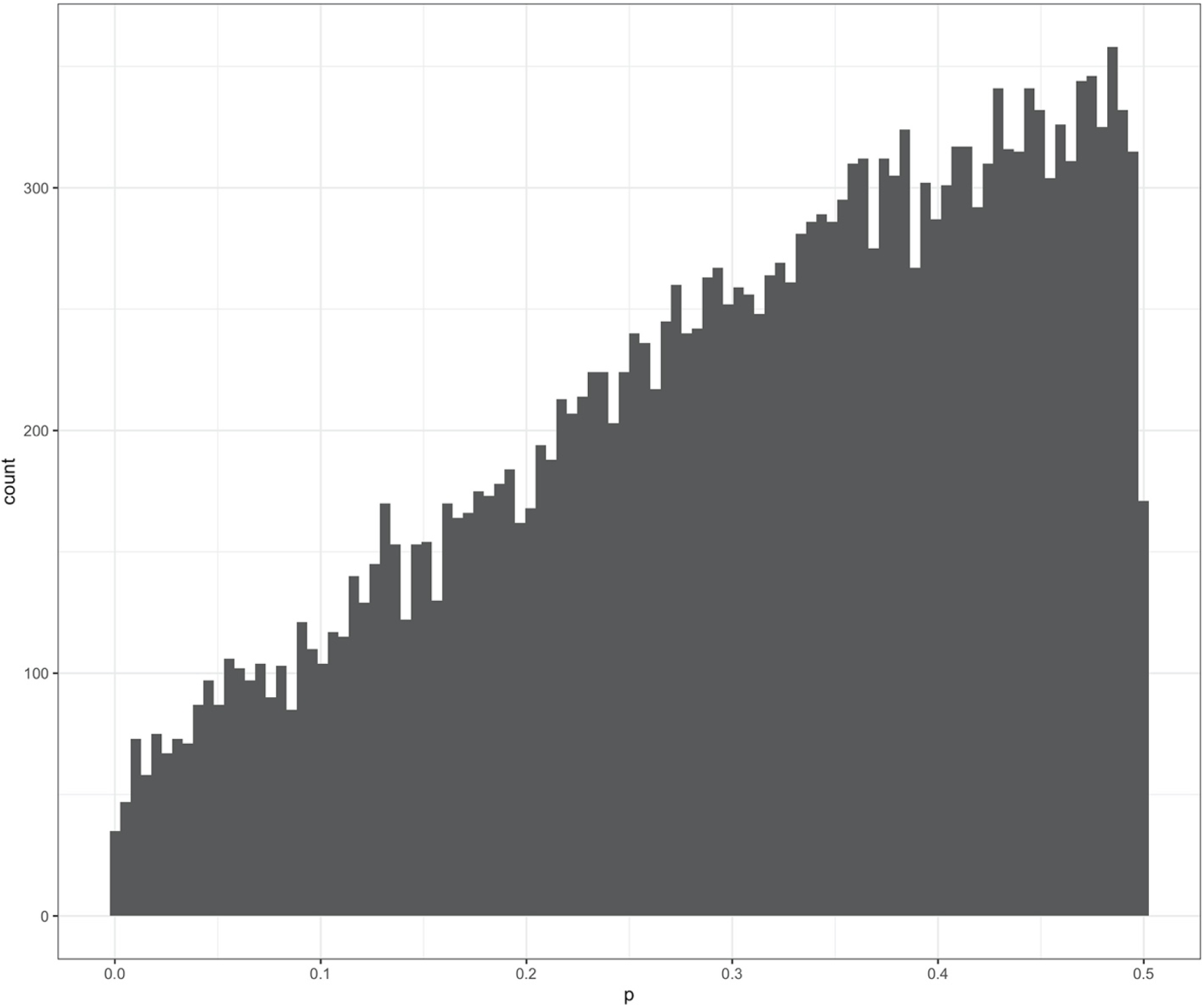
Un-adjusted p-values for all SNPs of each populations analysed with *spatpg* (11).

**Figure S6.**
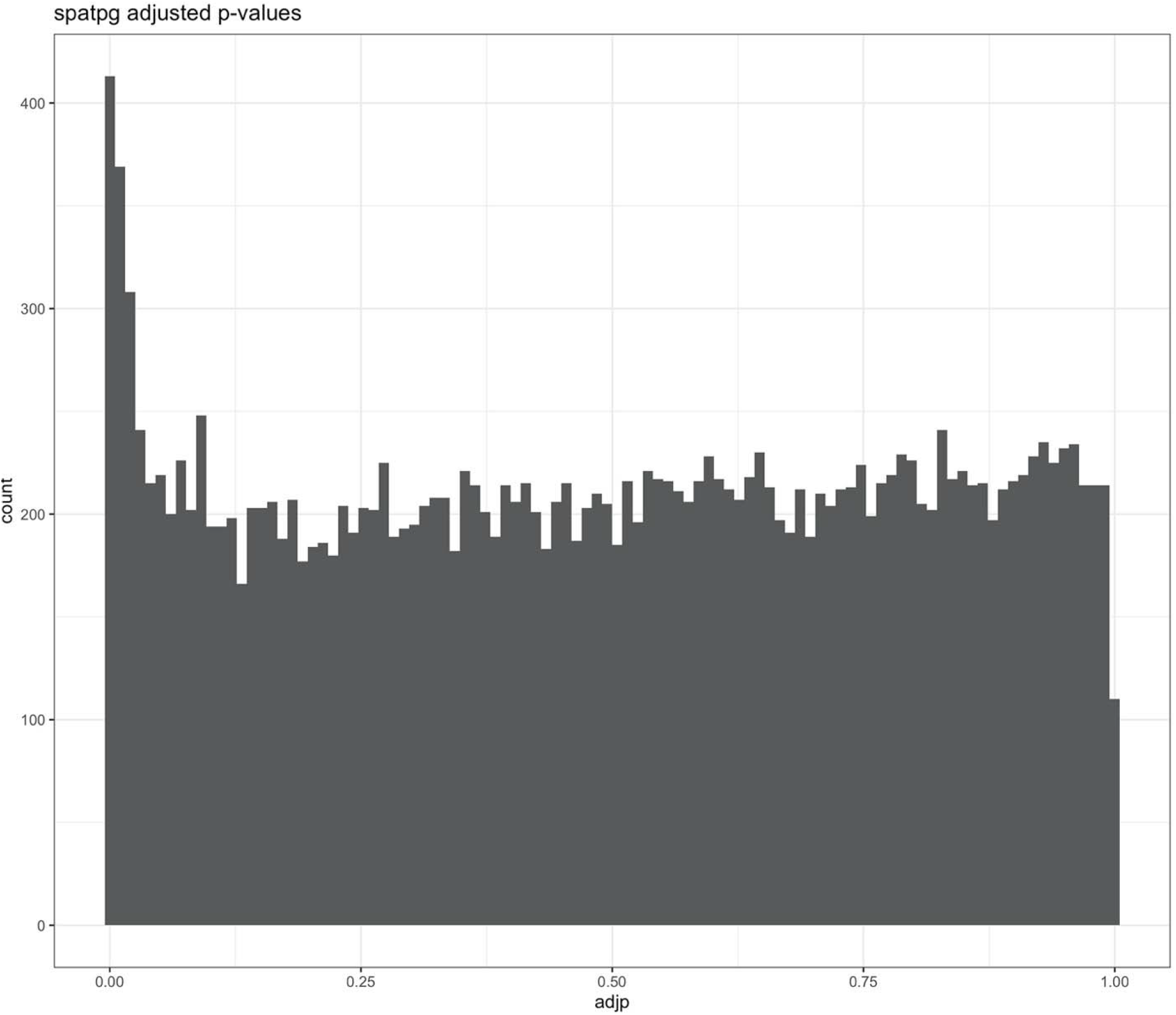
Adjusted p-values (12) for all SNPs of each population analysed with *spatpg* (10, 11).

**Figure S7.**
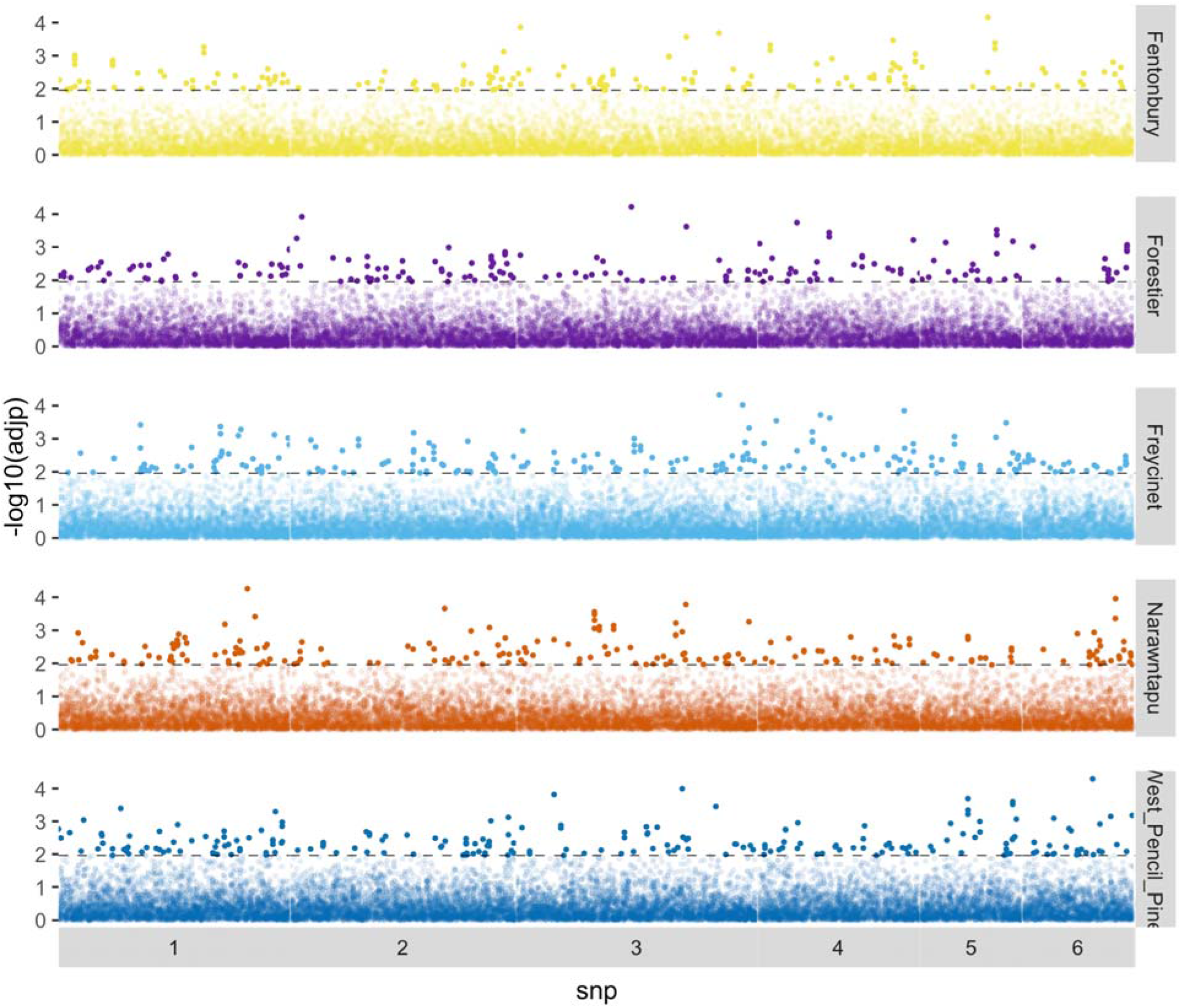
Allele frequency change (Δ*af*) for each population separately. SNPs in the top 1% are indicated by more opaque points. The threshold line for the top 1% within each population is indicated by a dashed line.

**Figure S8.**
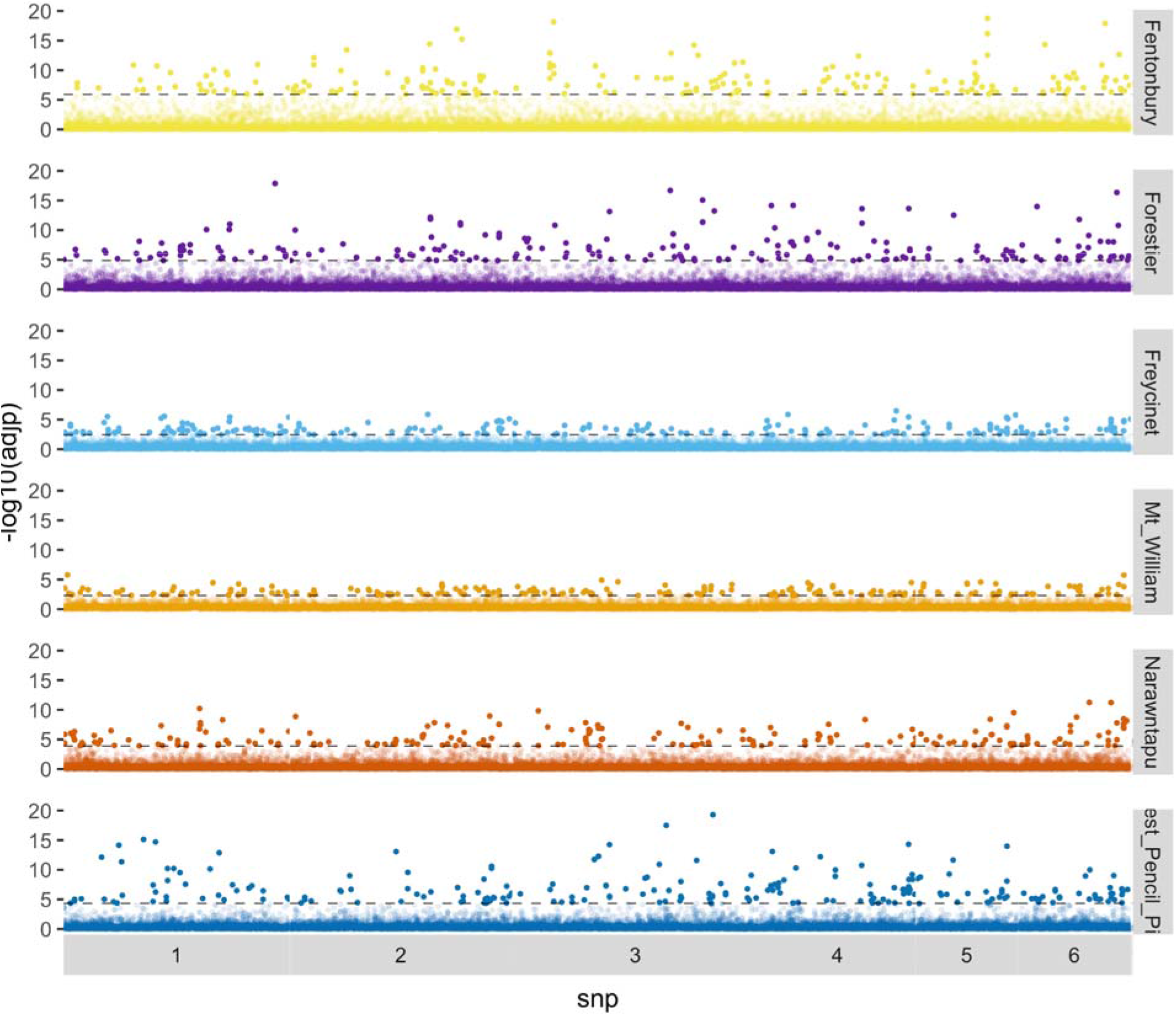
Signatures of selection as detected by *mm* for each population separately. SNPs in the top 1% are indicated by more opaque points. The threshold line for the top 1% within each population is indicated by a dashed line.

**Figure S9.**
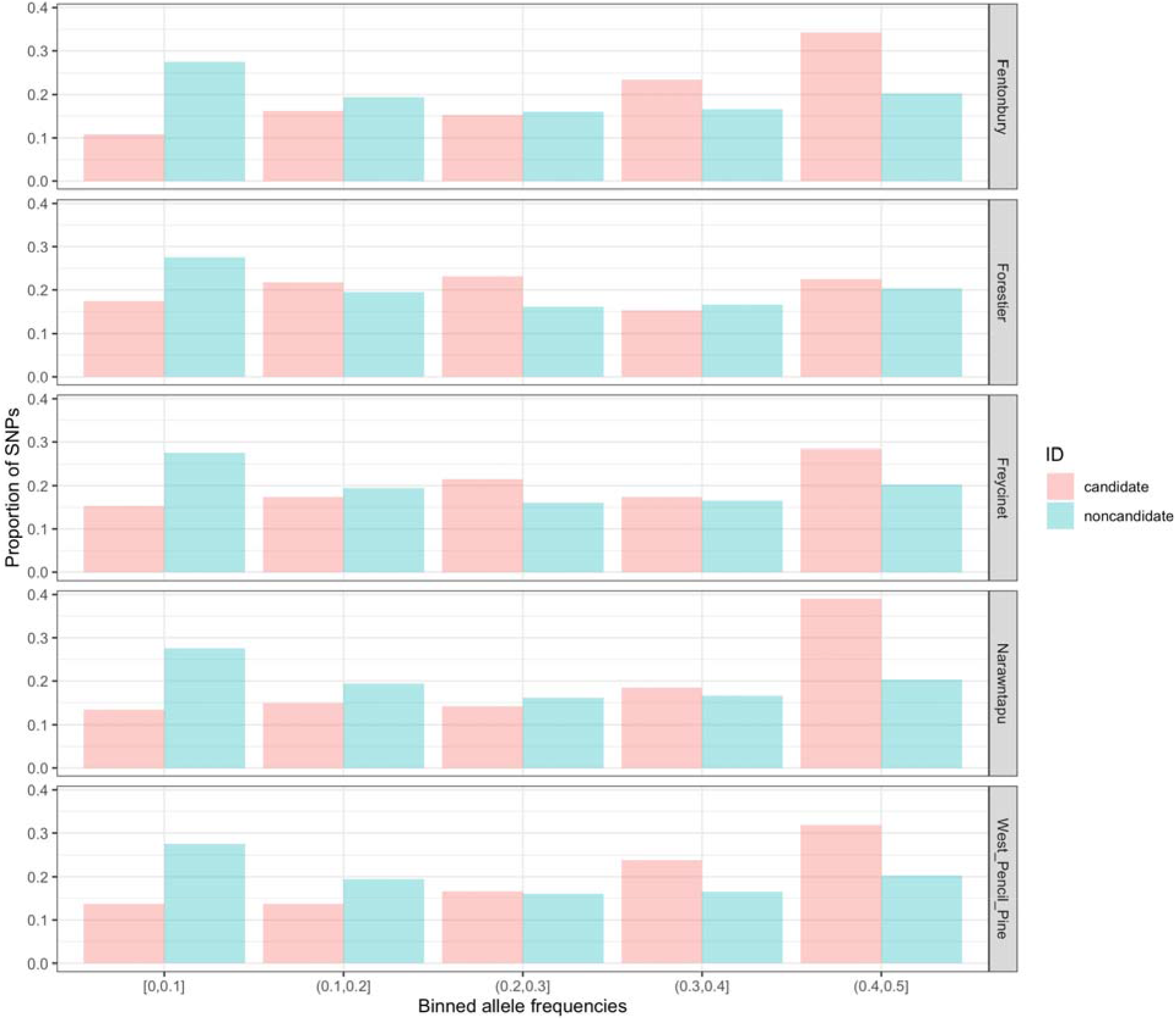
Binned initial allele frequency distributions for DCMS candidates and non-candidates across all populations with samples before DFTD became prevalent.

**Supplemental Figure S10.**
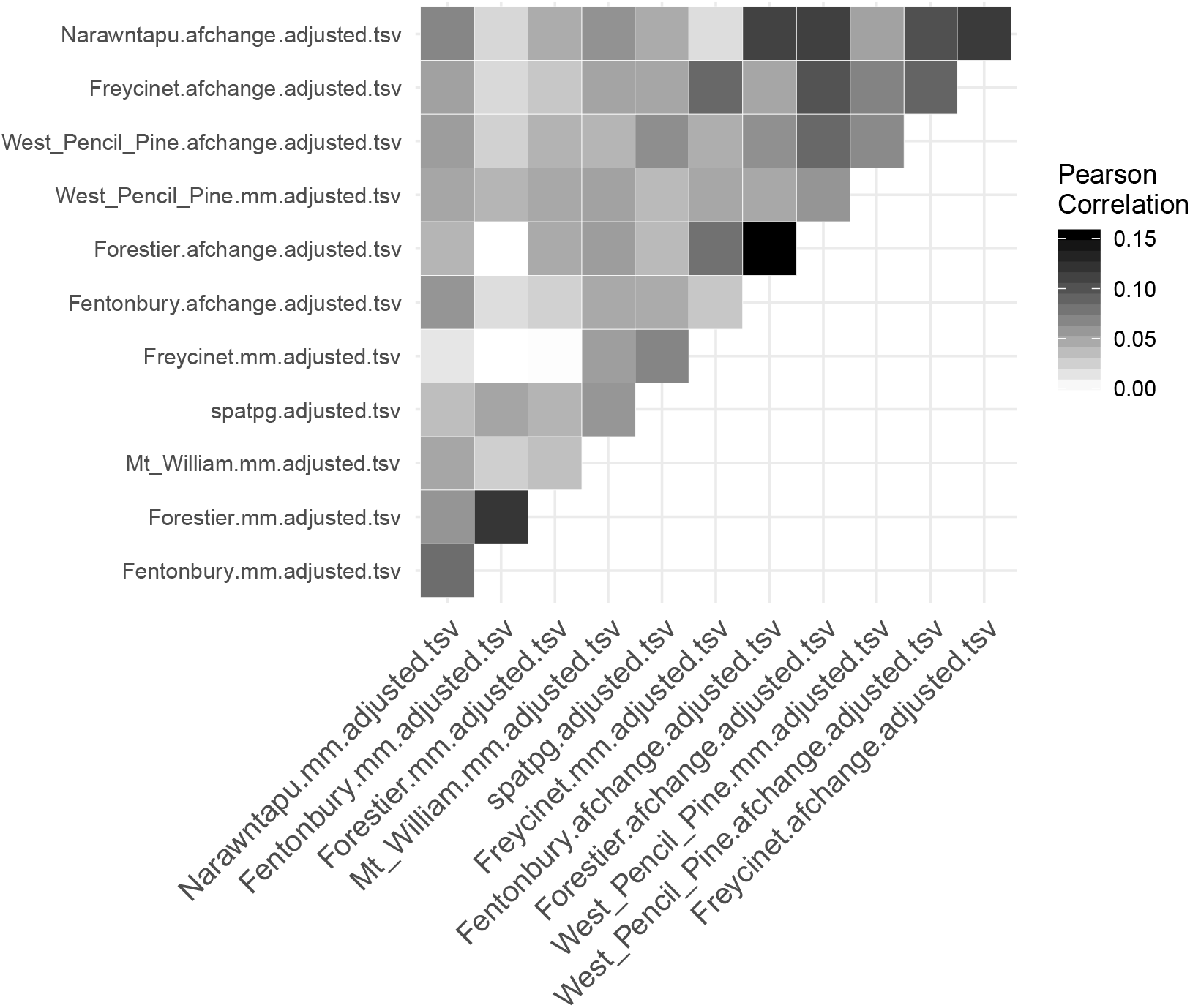
Correlations among elementary statistics: afchange = allele frequency change; mm = Mathieson and McVean (10), spatpg (11). Correlations are clustered by similarity along the x-axis.

**Figure S11.**
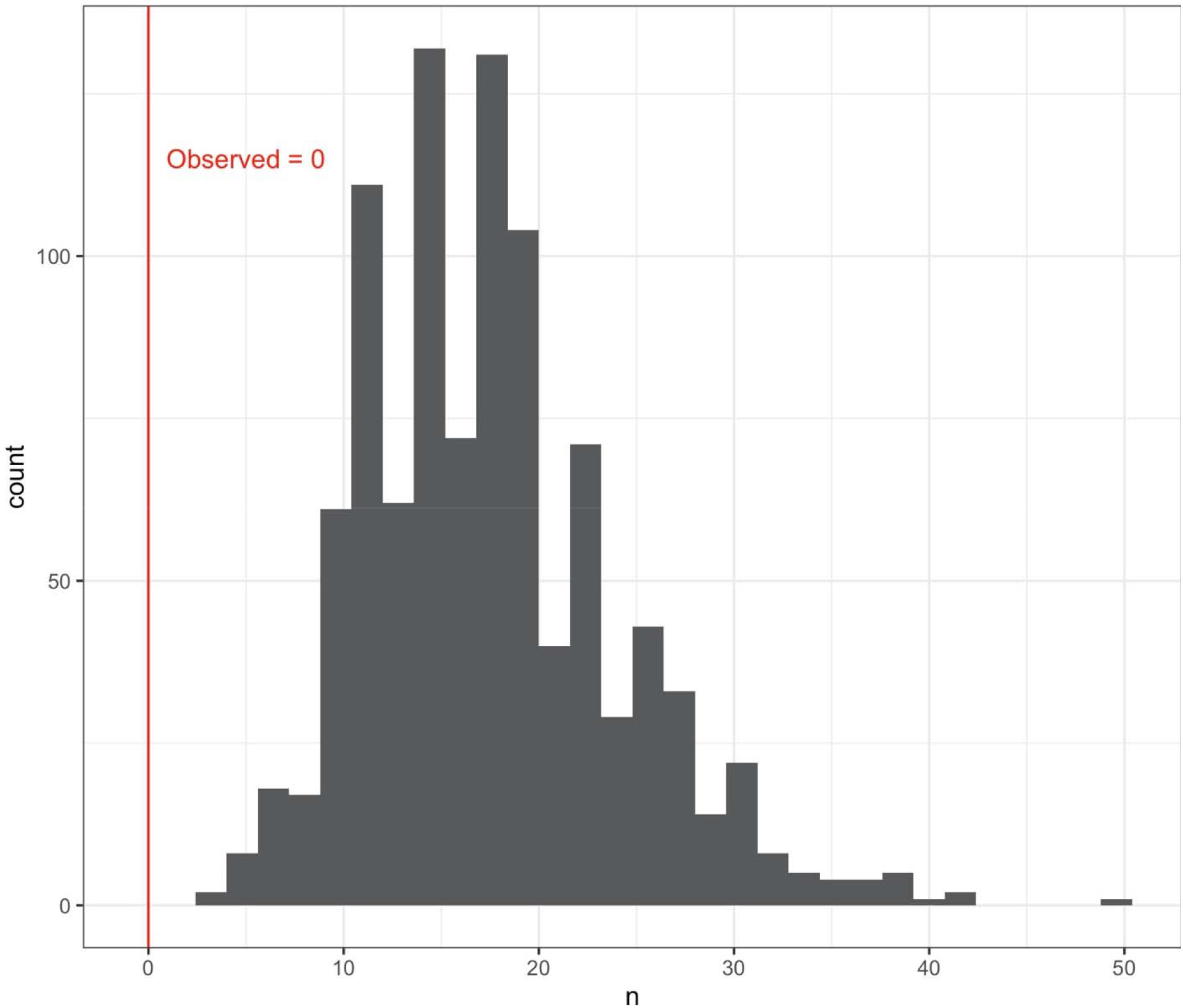
Histogram of shared MSigDB (35) gene set overlaps between contemporary and historical candidates, i.e. the signature of ongoing selection, in a permutation test of 1000 draws with replacement. We observed no shared gene set overlaps between contemporary and historical candidates. Of the 1000 permutations, 100% had fewer overlapping sets than observed.

**Supplemental Table S1.** Number and year of sampling across seven localities.

| Population | Year | Number of individuals |
| --- | --- | --- |
| wukalina/Mt. William | 2004 | 11 |
| wukalina/Mt. William | 2005 | 28 |
| wukalina/Mt. William | 2006 | 20 |
| wukalina/Mt. William | 2007 | 21 |
| wukalina/Mt. William | 2008 | 17 |
| wukalina/Mt. William | 2009 | 14 |
| wukalina/Mt. William | 2010 | 8 |
| wukalina/Mt. William | 2011 | 15 |
| wukalina/Mt. William | 2012 | 4 |
| wukalina/Mt. William | 2013 | 10 |
| wukalina/Mt. William | 2014 | 8 |
| **Freycinet** | 1999 | 107 |
| **Freycinet** | 2000 | 122 |
| **Freycinet** | 2001 | 71 |
| **Freycinet** | 2002 | 65 |
| **Freycinet** | 2003 | 38 |
| **Freycinet** | 2004 | 60 |
| **Freycinet** | 2005 | 54 |
| **Freycinet** | 2006 | 27 |
| **Freycinet** | 2007 | 32 |
| **Freycinet** | 2008 | 21 |
| **Freycinet** | 2009 | 7 |
| **Freycinet** | 2010 | 10 |
| **Freycinet** | 2011 | 10 |
| **Freycinet** | 2012 | 18 |
| **Freycinet** | 2013 | 12 |
| **Freycinet** | 2014 | 26 |
| Forestier | 2004 | 131 |
| Forestier | 2005 | 46 |
| Forestier | 2006 | 168 |
| Forestier | 2007 | 98 |
| Forestier | 2008 | 93 |
| Forestier | 2009 | 107 |
| Forestier | 2010 | 13 |
| Forestier | 2012 | 26 |
| Forestier | 2013 | 1 |
| Fentonbury | 2004 | 47 |
| Fentonbury | 2005 | 52 |
| Fentonbury | 2006 | 58 |
| Fentonbury | 2007 | 63 |
| Fentonbury | 2008 | 37 |
| Fentonbury | 2009 | 11 |
| **West Pencil Pine** | 2006 | 52 |
| **West Pencil Pine** | 2007 | 71 |
| **West Pencil Pine** | 2008 | 33 |
| **West Pencil Pine** | 2009 | 57 |
| **West Pencil Pine** | 2010 | 33 |
| **West Pencil Pine** | 2011 | 84 |
| **West Pencil Pine** | 2012 | 62 |
| **West Pencil Pine** | 2013 | 43 |
| **West Pencil Pine** | 2014 | 13 |
| Narawntapu | 1999 | 33 |
| Narawntapu | 2003 | 9 |
| Narawntapu | 2004 | 46 |
| Narawntapu | 2005 | 27 |
| Narawntapu | 2006 | 64 |
| Narawntapu | 2007 | 45 |
| Narawntapu | 2008 | 63 |
| Narawntapu | 2009 | 30 |
| Narawntapu | 2010 | 27 |
| Narawntapu | 2011 | 15 |
| Narawntapu | 2012 | 22 |

**Supplemental Table S2.** ANGSD (8) genotype calling settings.

| Option | Setting | Description |
| --- | --- | --- |
| -minMapQ | 40 | Minimum read mapping quality |
| -minQ | 25 | Minimum base Phred score |
| -baq | 2 | Perform extended base quality adjustment around indels (Li 2010) |
| -GL | 2 | Use the (old) GATK genotype likelihood model |
| -doMaf | 2 | Estimate allele frequencies assuming one known allele |
| -doMajorMinor | 4 | Use the reference allele as the known allele |

**Supplemental Table S3.**
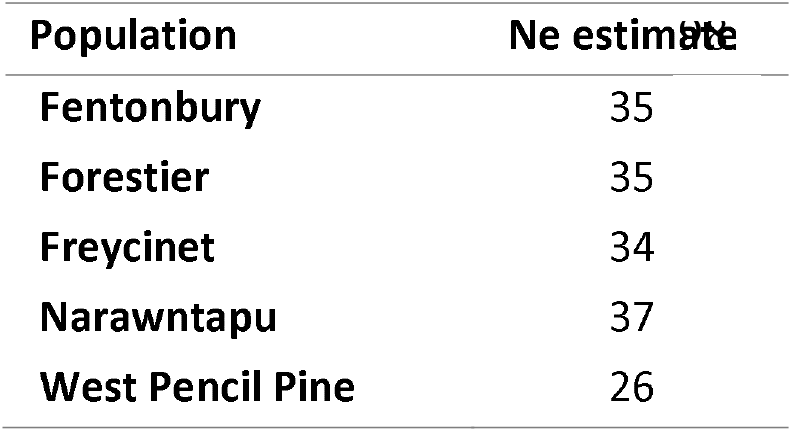
Estimates of effective population size.

**Supplemental Table S4.**
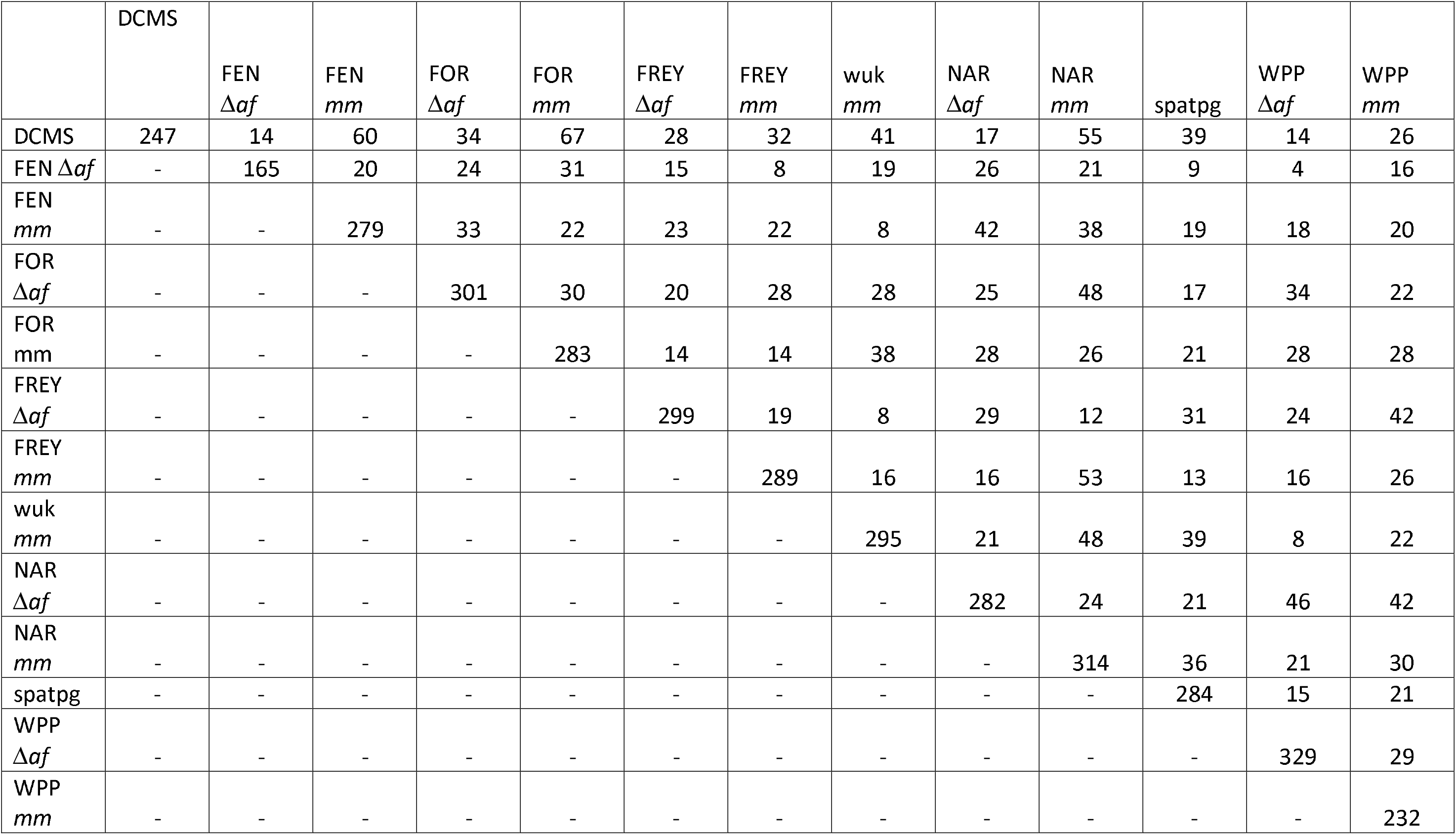
Number of shared genes within 10000 bp of the top 1% of SNPs for the DCMS list of contemporary candidates and each intermediate test. The total number of genes in each list is found in the diagonal. Populations are abbreviated as follows: FEN = Fentonbury, FOR = Forestier, FREY = Freycinet, wuk = wukalina/Mt. William, NAR= Narawntapu, and WPP = West Pencil Pine.

Supplemental Table S5. Annotated Tasmanian devil gene IDs of within 1000 bp of SNPs in the top 1% of *DCMS* scores; i.e. candidates for contemporary selection. Also provided at: https://github.com/Astahlke/contemporary_historical_sel_devils/blob/master/contemporary/angsd_2019-01-18/next/composite_stat/2019-2-22/results/annotation_top1.0/composite.snps.everything.top.genes.100000bp.txt.

Supplemental Table S6. Also provided at: https://github.com/Astahlke/contemporary_historical_sel_devils/blob/historical/sig_paml_results_21-1-13.csv. All PAML (16, 17) branch site test results for the 1,773 genes with inferred historical positive selection. Columns indicate: ensemble_gene_id = Ensembl gene ID for Tasmanian devils; likelihood = likelihood of the three-ratio model; p0 = proportion of sites belonging to site class 0, f0= dN/dS estimates for devil branch in site class 0 ; p1= proportion of sites in site class 1; f1= dN/dS estimates for devil branch in site class 0; p2 = proportion of sites belonging to site class 2a; f2 = dN/dS estimates for devil branch in site class 2a; p3 = proportion of sites belonging to site class 2b; dN/dS estimates for devil branch in site class 2b; f3 = dN/dS estimates for devil branch in site class 2b FDR = likelihood-ratio test p-values adjusted for multiple testing (38); p2a_2b = proportion of sites belonging to either cite class 2a or 2b

Site Class 0 = Background: 0< dN/dS<1; Foreground: 0< dN/dS<1
Site Class 1 = Background: dN/dS=1; Foreground: dN/dS=1
Site Class 2a = Background: dN/dS<1; Foreground: dN/dS≥1
Site Class 2b = Background: dN/dS=1; Foreground: dN/dS≥1

Supplemental Table S7. Sixteen candidate genes for both historical positive selection (p < 0.05 in PAML branch-site test) and a response to contemporary selection from DFTD (top 1% DCMS scores). Novel genes are annotated here by the Ensembl Compara gene family pipeline (15). * denotes genes previously associated with disease-related phenotypes in many of the same devils (32).

## References

1. Barreiro LB, Quintana-Murci L. From evolutionary genetics to human immunology: how selection shapes host defence genes. Nat Rev Genet. 2010;11(1):17–30.

2. Epstein B, Jones M, Hamede R, Hendricks S, McCallum H, Murchison EP, et al. Rapid evolutionary response to a transmissible cancer in Tasmanian devils. Nature communications. 2016;7:12684.

3. De Castro F, Bolker B. Mechanisms of disease◻induced extinction. Ecology Letters. 2005;8(1):117–26.

4. Smith KF, Sax DF, Lafferty KD. Evidence for the role of infectious disease in species extinction and endangerment. Conservation biology. 2006;20(5):1349–57.

5. Jones KE, Patel NG, Levy MA, Storeygard A, Balk D, Gittleman JL, et al. Global trends in emerging infectious diseases. Nature. 2008;451(7181):990–3.

6. Lafferty KD. The ecology of climate change and infectious diseases. Ecology. 2009;90(4):888–900.

7. McCallum H, Dobson A. Detecting disease and parasite threats to endangered species and ecosystems. Trends in ecology & evolution. 1995;10(5):190–4.

8. Storfer A, Kozakiewicz CP, Beer MA, Savage AE. Applications of Population Genomics for Understanding and Mitigating Wildlife Disease. 2020.

9. McKnight DT, Schwarzkopf L, Alford RA, Bower DS, Zenger KR. Effects of emerging infectious diseases on host population genetics: a review. Conservation genetics. 2017;18(6):1235–45.

10. Brandies P, Peel E, Hogg CJ, Belov K. The Value of Reference Genomes in the Conservation of Threatened Species. Genes. 2019;10(11):846.

11. Li W-H. Unbiased estimation of the rates of synonymous and nonsynonymous substitution. Journal of molecular evolution. 1993;36(1):96–9.

12. Yang Z. Likelihood ratio tests for detecting positive selection and application to primate lysozyme evolution. Molecular biology and evolution. 1998;15(5):568–73.

13. Shultz AJ, Sackton TB. Immune genes are hotspots of shared positive selection across birds and mammals. Elife. 2019;8:e41815.

14. Mathieson I, McVean G. Estimating selection coefficients in spatially structured populations from time series data of allele frequencies. Genetics. 2013;193(3):973–84.

15. Gompert Z. Bayesian inference of selection in a heterogeneous environment from genetic time-series data. Mol Ecol. 2016;25(1):121–34.

16. Yeaman S, Gerstein AC, Hodgins KA, Whitlock MC. Quantifying how constraints limit the diversity of viable routes to adaptation. PLoS genetics. 2018;14(10):e1007717.

17. Baird NA, Etter PD, Atwood TS, Currey MC, Shiver AL, Lewis ZA, et al. Rapid SNP discovery and genetic mapping using sequenced RAD markers. PloS one. 2008;3(10):e3376.

18. Andrews KR, Good JM, Miller MR, Luikart G, Hohenlohe PA. Harnessing the power of RADseq for ecological and evolutionary genomics. Nature Reviews Genetics. 2016;17(2):81.

19. Hawkins CE, Baars C, Hesterman H, Hocking GJ, Jones ME, Lazenby B, et al. Emerging disease and population decline of an island endemic, the Tasmanian devil Sarcophilus harrisii. Biological Conservation. 2006;131(2):307–24.

20. Pearse AM, Swift K. Allograft theory: transmission of devil facial-tumour disease. Nature. 2006;439(7076):549.

21. Hamilton DG, Jones ME, Cameron EZ, McCallum H, Storfer A, Hohenlohe PA, et al. Rate of intersexual interactions affects injury likelihood in Tasmanian devil contact networks. Behavioral Ecology. 2019;30(4):1087–95.

22. Hamede RK, McCallum H, Jones M. Seasonal, demographic and density◻related patterns of contact between Tasmanian devils (Sarcophilus harrisii): Implications for transmission of devil facial tumour disease. Austral Ecology. 2008;33(5):614–22.

23. Pye R, Hamede R, Siddle HV, Caldwell A, Knowles GW, Swift K, et al. Demonstration of immune responses against devil facial tumour disease in wild Tasmanian devils. Biol Lett. 2016;12(10).

24. Siddle HV, Kreiss A, Tovar C, Yuen CK, Cheng Y, Belov K, et al. Reversible epigenetic down-regulation of MHC molecules by devil facial tumour disease illustrates immune escape by a contagious cancer. Proceedings of the National Academy of Sciences. 2013;110(13):5103–8.

25. Murchison EP, Tovar C, Hsu A, Bender HS, Kheradpour P, Rebbeck CA, et al. The Tasmanian devil transcriptome reveals Schwann cell origins of a clonally transmissible cancer. Science. 2010;327(5961):84–7.

26. Lazenby BT, Tobler MW, Brown WE, Hawkins CE, Hocking GJ, Hume F, et al. Density trends and demographic signals uncover the long◻term impact of transmissible cancer in Tasmanian devils. Journal of Applied Ecology. 2018;55(3):1368–79.

27. McCallum H, Jones M, Hawkins C, Hamede R, Lachish S, Sinn DL, et al. Transmission dynamics of Tasmanian devil facial tumor disease may lead to disease◻induced extinction. Ecology. 2009;90(12):3379–92.

28. Hubert J-N, Zerjal T, Hospital F. Cancer-and behavior-related genes are targeted by selection in the Tasmanian devil (Sarcophilus harrisii). PloS one. 2018;13(8):e0201838.

29. Fraik AK, Margres MJ, Epstein B, Barbosa S, Jones M, Hendricks S, et al. Disease swamps molecular signatures of genetic-environmental associations to abiotic factors in Tasmanian devil (Sarcophilus harrisii) populations. Evolution. 2020;n/a(n/a).

30. Wells K, Hamede RK, Jones ME, Hohenlohe PA, Storfer A, McCallum HI. Individual and temporal variation in pathogen load predicts long◻term impacts of an emerging infectious disease. Ecology. 2019;100(3):e02613.

31. Brüniche–Olsen A, Jones ME, Burridge CP, Murchison EP, Holland BR, Austin JJ. Ancient DNA tracks the mainland extinction and island survival of the Tasmanian devil. Journal of biogeography. 2018;45(5):963–76.

32. Brüniche-Olsen A, Jones ME, Austin JJ, Burridge CP, Holland BR. Extensive population decline in the Tasmanian devil predates European settlement and devil facial tumour disease. Biology letters. 2014;10(11):20140619.

33. Patton AH, Margres MJ, Stahlke AR, Hendricks S, Lewallen K, Hamede RK, et al. Contemporary demographic reconstruction methods are robust to genome assembly quality: A case study in Tasmanian Devils. Molecular biology and evolution. 2019;36(12):2906–21.

34. Siddle HV, Kreiss A, Eldridge MDB, Noonan E, Clarke CJ, Pyecroft S, et al. Transmission of a fatal clonal tumor by biting occurs due to depleted MHC diversity in a threatened carnivorous marsupial. Proceedings of the National Academy of Sciences. 2007;104(41):16221–6.

35. Pye RJ, Pemberton D, Tovar C, Tubio JMC, Dun KA, Fox S, et al. A second transmissible cancer in Tasmanian devils. Proceedings of the National Academy of Sciences. 2016;113(2):374–9.

36. James S, Jennings G, Kwon YM, Stammnitz M, Fraik A, Storfer A, et al. Tracing the rise of malignant cell lines: Distribution, epidemiology and evolutionary interactions of two transmissible cancers in Tasmanian devils. Evolutionary applications. 2019;12(9):1772–80.

37. Stammnitz MR, Coorens THH, Gori KC, Hayes D, Fu B, Wang J, et al. The origins and vulnerabilities of two transmissible cancers in Tasmanian devils. Cancer Cell. 2018;33(4):607–19.

38. Deakin JE, Bender HS, Pearse A-M, Rens W, O’Brien PCM, Ferguson-Smith MA, et al. Genomic restructuring in the Tasmanian devil facial tumour: chromosome painting and gene mapping provide clues to evolution of a transmissible tumour. PLoS genetics. 2012;8(2):e1002483.

39. Peck SJ, Michael SA, Knowles G, Davis A, Pemberton D. Causes of mortality and severe morbidity requiring euthanasia in captive Tasmanian devils (Sarcophilus harrisii) in Tasmania. Australian veterinary journal. 2019;97(4):89–92.

40. Hamede RK, McCallum H, Jones M. Biting injuries and transmission of T asmanian devil facial tumour disease. Journal of Animal Ecology. 2013;82(1):182–90.

41. Patchett AL, Flies AS, Lyons AB, Woods GM. Curse of the devil: molecular insights into the emergence of transmissible cancers in the Tasmanian devil (Sarcophilus harrisii). Cellular and Molecular Life Sciences. 2020.

42. Hogg CJ, Lee AV, Srb C, Hibbard C. Metapopulation management of an Endangered species with limited genetic diversity in the presence of disease: the Tasmanian devil Sarcophilus harrisii. International Zoo Yearbook. 2017;51(1):137–53.

43. Hohenlohe PA, McCallum HI, Jones ME, Lawrance MF, Hamede RK, Storfer A. Conserving adaptive potential: lessons from Tasmanian devils and their transmissible cancer. Conservation Genetics. 2019;20(1):81–7.

44. Wright BR, Farquharson KA, McLennan EA, Belov K, Hogg CJ, Grueber CE. A demonstration of conservation genomics for threatened species management. Molecular Ecology Resources. 2020.

45. Grueber CE, Peel E, Wright B, Hogg CJ, Belov K. A Tasmanian devil breeding program to support wild recovery. Reproduction, Fertility and Development. 2019;31(7):1296–304.

46. Ali OA, O’Rourke SM, Amish SJ, Meek MH, Luikart G, Jeffres C, et al. RAD Capture (Rapture): Flexible and efficient sequence-based genotyping. Genetics. 2016;202(2):389–400.

47. Margres MJ, Jones ME, Epstein B, Kerlin DH, Comte S, Fox S, et al. Large◻effect loci affect survival in Tasmanian devils (Sarcophilus harrisii) infected with a transmissible cancer. Molecular ecology. 2018;27(21):4189–99.

48. Hamede RK, Pearse AM, Swift K, Barmuta LA, Murchison EP, Jones ME. Transmissible cancer in Tasmanian devils: localized lineage replacement and host population response. Proc Biol Sci. 2015;282(1814).

49. Ma Y, Ding X, Qanbari S, Weigend S, Zhang Q, Simianer H. Properties of different selection signature statistics and a new strategy for combining them. Heredity. 2015;115(5):426.

50. Jorde PE, Ryman N. Unbiased estimator for genetic drift and effective population size. Genetics. 2007;177(2):927–35.

51. Do C, Waples RS, Peel D, Macbeth GM, Tillett BJ, Ovenden JR. NeEstimator v2: re◻implementation of software for the estimation of contemporary effective population size (Ne) from genetic data. Molecular ecology resources. 2014;14(1):209–14.

52. Kelly JK, Hughes KA. Pervasive linked selection and intermediate-frequency alleles are implicated in an evolve-and-resequencing experiment of Drosophila simulans. Genetics. 2019;211(3):943–61.

53. Mikkelsen TS, Wakefield MJ, Aken B, Amemiya CT, Chang JL, Duke S, et al. Genome of the marsupial Monodelphis domestica reveals innovation in non-coding sequences. Nature. 2007;447(7141):167–77.

54. Renfree MB, Papenfuss AT, Deakin JE, Lindsay J, Heider T, Belov K, et al. Genome sequence of an Australian kangaroo, Macropus eugenii, provides insight into the evolution of mammalian reproduction and development. Genome biology. 2011;12(8):1–26.

55. Aken BL, Achuthan P, Akanni W, Amode MR, Bernsdorff F, Bhai J, et al. Ensembl 2017. Nucleic acids research. 2016;45(D1):D635–D42.

56. Johnson RN, O’Meally D, Chen Z, Etherington GJ, Ho SYW, Nash WJ, et al. Adaptation and conservation insights from the koala genome. Nature genetics. 2018;50(8):1102.

57. Yang Z. PAML 4: phylogenetic analysis by maximum likelihood. Molecular biology and evolution. 2007;24(8):1586–91.

58. Yang Z. PAML: a program package for phylogenetic analysis by maximum likelihood. Bioinformatics. 1997;13(5):555–6.

59. Mitchell KJ, Pratt RC, Watson LN, Gibb GC, Llamas B, Kasper M, et al. Molecular phylogeny, biogeography, and habitat preference evolution of marsupials. Molecular biology and evolution. 2014;31(9):2322–30.

60. Anisimova M, Bielawski JP, Yang Z. Accuracy and power of the likelihood ratio test in detecting adaptive molecular evolution. Molecular biology and evolution. 2001;18(8):1585–92.

61. Bielawski JP, Yang Z. Maximum likelihood methods for detecting adaptive protein evolution. Statistical methods in molecular evolution: Springer; 2005. p. 103–24.

62. Shaffer JP. Multiple hypothesis testing. Annual review of psychology. 1995;46(1):561–84.

63. Massey Jr FJ. The Kolmogorov-Smirnov test for goodness of fit. Journal of the American statistical Association. 1951;46(253):68–78.

64. Scholz FW, Stephens MA. K-sample Anderson–Darling tests. Journal of the American Statistical Association. 1987;82(399):918–24.

65. Szkiba D, Kapun M, von Haeseler A, Gallach M. SNP2GO: functional analysis of genome-wide association studies. Genetics. 2014;197(1):285–9.

66. Mi H, Muruganujan A, Ebert D, Huang X, Thomas PD. PANTHER version 14: more genomes, a new PANTHER GO-slim and improvements in enrichment analysis tools. Nucleic acids research. 2018;47(D1):D419–D26.

67. Subramanian A, Tamayo P, Mootha VK, Mukherjee S, Ebert BL, Gillette MA, et al. Gene set enrichment analysis: a knowledge-based approach for interpreting genome-wide expression profiles. Proceedings of the National Academy of Sciences. 2005;102(43):15545–50.

68. Wright B, Willet CE, Hamede R, Jones M, Belov K, Wade CM. Variants in the host genome may inhibit tumour growth in devil facial tumours: evidence from genome-wide association. Sci Rep. 2017;7(1):423.

69. Wu K, Li Z, Cai S, Tian L, Chen K, Wang J, et al. EYA1 phosphatase function is essential to drive breast cancer cell proliferation through cyclin D1. Cancer research. 2013;73(14):4488–99.

70. Cai S, Cheng X, Liu Y, Lin Z, Zeng W, Yang C, et al. EYA1 promotes tumor angiogenesis by activating the PI3K pathway in colorectal cancer. Experimental cell research. 2018;367(1):37–46.

71. Meurs EF, Galabru J, Barber GN, Katze MG, Hovanessian AG. Tumor suppressor function of the interferon-induced double-stranded RNA-activated protein kinase. Proceedings of the National Academy of Sciences. 1993;90(1):232–6.

72. Brüniche-Olsen A, Austin JJ, Jones ME, Holland BR, Burridge CP. Detecting selection on temporal and spatial scales: a genomic time-series assessment of selective responses to devil facial tumor disease. PloS one. 2016;11(3).

73. Hendricks S, Epstein B, Schönfeld B, Wiench C, Hamede R, Jones M, et al. Conservation implications of limited genetic diversity and population structure in Tasmanian devils (Sarcophilus harrisii). Conservation Genetics. 2017;18(4):977–82.

74. Margres MJ, Ruiz-Aravena M, Hamede R, Jones ME, Lawrance MF, Hendricks SA, et al. The genomic basis of tumor regression in Tasmanian devils (Sarcophilus harrisii). Genome biology and evolution. 2018;10(11):3012–25.

75. Fang WH, Wang Q, Li HM, Ahmed M, Kumar P, Kumar S. PAX 3 in neuroblastoma: oncogenic potential, chemosensitivity and signalling pathways. Journal of cellular and molecular medicine. 2014;18(1):38–48.

76. Zhang J. Frequent false detection of positive selection by the likelihood method with branch-site models. Molecular Biology and Evolution. 2004;21(7):1332–9.

77. Kosiol C, Vinař T, da Fonseca RR, Hubisz MJ, Bustamante CD, Nielsen R, et al. Patterns of positive selection in six mammalian genomes. PLoS genetics. 2008;4(8).

78. Nelson WJ, Nusse R. Convergence of Wnt, ß-catenin, and cadherin pathways. Science. 2004;303(5663):1483–7.

79. Metzger MJ, Reinisch C, Sherry J, Goff SP. Horizontal transmission of clonal cancer cells causes leukemia in soft-shell clams. Cell. 2015;161(2):255–63.

80. Strakova A, Murchison EP. The cancer which survived: insights from the genome of an 11000 year-old cancer. Curr Opin Genet Dev. 2015;30:49–55.

81. Casanova J-L, Abel L, editors. Human genetics of infectious diseases: Unique insights into immunological redundancy2018 2018: Elsevier.

82. Guiler E. The Tasmanian devil: St. David’s Park Pub.; 1992.

83. Miller IF, Metcalf CJE. Evolving resistance to pathogens. Science. 2019;363(6433):1277–8.

84. Gandon S, Michalakis Y. Evolution of parasite virulence against qualitative or quantitative host resistance. Proceedings of the Royal Society of London Series B: Biological Sciences. 2000;267(1447):985–90.

85. Patton AH, Lawrance MF, Margres MJ, Kozakiewicz CP, Hamede R, Ruiz-Aravena M, et al. A transmissible cancer shifts from emergence to endemism in Tasmanian devils. Science. 2020;370(6522).

86. Kardos M, Shafer ABA. The Peril of Gene-Targeted Conservation. Trends Ecol Evol. 2018;33(11):827–39.

87. McLennan EA, Grueber CE, Wise P, Belov K, Hogg CJ. Mixing genetically differentiated populations successfully boosts diversity of an endangered carnivore. Animal Conservation. 2020.

88. Wright B, Morris K, Grueber CE, Willet CE, Gooley R, Hogg CJ, et al. Development of a SNP-based assay for measuring genetic diversity in the Tasmanian devil insurance population. BMC Genomics. 2015;16:791.

89. Campbell NR, Harmon SA, Narum SR. Genotyping-in-Thousands by sequencing (GT-seq): A cost effective SNP genotyping method based on custom amplicon sequencing. Mol Ecol Resour. 2015;15(4):855–67.

90. von Thaden A, Nowak C, Tiesmeyer A, Reiners TE, Alves PC, Lyons LA, et al. Applying genomic data in wildlife monitoring: Development guidelines for genotyping degraded samples with reduced single nucleotide polymorphism panels. Molecular Ecology Resources. 2020;20(3).

91. Margres MJ, Ruiz-Aravena M, Hamede R, Chawla K, Patton AH, Lawrance MF, et al. Spontaneous tumor regression in Tasmanian devils associated with RASL11A activation. Genetics. 2020.

